# Day-to-day spontaneous social behaviours is quantitatively and qualitatively affected in a 16p11.2 deletion mouse model

**DOI:** 10.1101/2022.09.16.508234

**Authors:** Anna Rusu, Claire Chevalier, Fabrice de Chaumont, Valérie Nalesso, Véronique Brault, Yann Hérault, Elodie Ey

## Abstract

**Background:** Autism spectrum disorders affect more than one percent of the population, impairing social communication and increasing stereotyped behaviours. A micro-deletion of the 16p11.2 BP4-BP5 chromosomic region has been identified in one percent of patients also displaying intellectual disabilities. In mouse models generated to understand the mechanisms of this deletion, learning and memory deficits were pervasive in most genetic backgrounds, while social communication deficits were only detected in some models.

**Methods:** Based on previous study (Arbogast et al. 2016 PLoS genetics), we itemized the social deficits in the mouse model of 16p11.2 deletion on a hybrid C57BL/6NxC3H.*Pde6b^+^* genetic background. We examined whether behavioural deficits were visible over long-term observation periods, to parallel everyday-life assessment of patients. We recorded the individual and social behaviours of 16p11.2 Del/+ mice and their wild-type littermates from both sexes in long-term (over two and three consecutive nights) social interactions of familiar mixed-genotype quartets of males and of females, and of same-genotype unfamiliar female pairs.

**Results:** We observed that Del/+ mice of both sexes increased significantly their locomotor activity compared to wild-type littermates over long-term monitoring. In the social domain, Del/+ mice of both sexes displayed widespread deficits over long-term monitoring, even more so in males than in females in quartets of familiar individuals. In pairs, significant perturbations of the organisation of the social communication and behaviours appeared in Del/+ females.

**Discussion:** Altogether, this suggests that, over long recording periods, the phenotype of the 16p11.2 Del/+ mice was differently affected in the locomotor activity and the social domains and between the two sexes. These findings confirm the importance of testing models in long-term conditions to provide a comprehensive view of their phenotype that will refine the study of cellular and molecular mechanisms and complement pre-clinical targeted therapeutic trials.

## Background

Autism spectrum disorder (ASD) is a neurodevelopmental condition characterised at the clinical level by atypical social interactions and communication, as well as stereotyped behaviours and restricted interests (American Psychiatric Association, 2013). This condition affects not only the patient but also his/her whole family. There exists a large range of severity between patients, who can also present severe comorbidities such as intellectual disability (ID), epilepsy, sleep disorders or hyper/hypo-sensitivity (Matson and Goldin, 2013). The prevalence of ASD is more than 1% of the general population with more males than females (Fombonne et al., 2021; Maenner et al., 2021). Potential causes can be environmental or genetic. Among these, copy number variations in the 16p11.2 region have been identified as one of the most frequent genetic causes of ASD (Weiss et al., 2008). This region of 600kb between two repeated sequences named BP4 and BP5 includes 28 genes that can be either deleted or duplicated ((Portmann et al., 2014): 550 kb and 26 genes; (Rein and Yan, 2020): 500-600 kb containing 27-29 genes). The duplication has been robustly linked with schizophrenia (McCarthy et al., 2009; Bergen et al., 2012), while the deletion is associated with 1% of ASD cases accompanied by ID (Jacquemont et al., 2011).

Patients with a deletion in the 16p11.2 region present diverse phenotypes, such as ASD (15% of cases), speech and language disorders (80-90% of cases; (Rosenfeld et al., 2010; Mei et al., 2018)), abnormal adaptive behaviours, cognitive behaviours and repetitive behaviours (at least one of these domains affected in 70-90% of cases; (Zufferey et al., 2012; Hanson et al., 2015)), sleep disorders (80% of cases; (Goldman et al., 2011)), ID (20% of cases; (Zufferey et al., 2012)), hyperactivity or attention disorder (30-40% of cases; (Rein and Yan, 2020)), developmental delay (100% of cases; (Rosenfeld et al., 2010; Rein and Yan, 2020)), epilepsy (10-20%; (Rosenfeld et al., 2010; Rein and Yan, 2020)), facial dysmorphia (>20% of cases; (Rosenfeld et al., 2010; Qiu et al., 2019; Rein and Yan, 2020)), obesity and macrocephaly (Zufferey et al., 2012). Patients may also present atypical brain anatomy, with abnormalities in the cerebellar tonsil (Steinman et al., 2016), auditory and speech pathways, as well as in the cortical and striatal structures (Maillard et al., 2015).

Overall and of particular interest as a phenotype related to ASD, these patients frequently display social interaction and communication impairments, especially in speech development (Benedetti et al., 2022). They also show poorer adaptative abilities in their daily life compared to controls (Hanson et al., 2015). These two aspects evaluated both during short-term clinical examination and during every-day life observation constitute keys points to examine in pre-clinical models of ASD to grab a more complete overview of their phenotype.

The homologous region of the 16p11.2 lies in mouse chromosome 7F3 (Horev et al., 2011; Portmann et al., 2014; Arbogast et al., 2016). Four mouse models were generated, differing in their genetic background and in the size of the deleted chromosomic region ((Horev et al., 2011; Portmann et al., 2014; Arbogast et al., 2016; Nakamura et al., 2021); see review in **Supplementary Table I** and in (Benedetti et al., 2022)). These models were further characterised either in the same genetic background or in different backgrounds. All of the four models displayed a reduced body weight compared to their wild-type littermates. Most of them displayed typical or increased activity in the short-term exploration of an open field, and increased activity over long-term (over one day or more) recordings compared to wild-type mice. Stereotyped behaviours remained subtle. Deficits in novel object recognition were recurrently highlighted in the different models. Sensory abilities were minimally affected, except in one model that appeared to be deaf due certainly to the genetic background (Portmann et al., 2014; Yang et al., 2015b). Given our interest on ASD-related phenotype, we focused our attention on social abilities. Over the different models, the variability of the social deficit attracted attention (Benedetti et al., 2022). One potential confounding factor is the genetic background of the models. Indeed, for a similar deletion and using identical test procedures, our laboratory did not detect robust social deficits in mutant mice generated on a pure C57BL/6N (B6N) background, while the same deletion crossed to obtain a F1 B6NC3B background (C3B for sighted C3H.*Pde6b^+^*; see the material and methods section) provoked a reduced time of sniffing in mutant mice compared to wild-type littermates (Arbogast et al., 2016). We chose therefore this last model generated on a mixed background to investigate further the everyday-life-deficits in the social and communication domains.

In the clinical practice, practitioners developed approaches focusing on the impairments in the everyday life of patients, with the Social Responsiveness Scale filled by caretakers and the Brief Observation of Social Communication Change (BOSCC) clinical evaluation made on videos of spontaneous play between parents/caretakers and patients (Grzadzinski et al., 2016). In the present study, we used a similar approach to grab a phenotype closer to the everyday life of animals. Therefore, we focused on spontaneous social interactions over long-term observation periods, dissecting the different types of body contacts and their dynamics (de Chaumont et al., 2019, 2021) to examine social orientation, seeking and maintenance of social contacts (Chevallier et al., 2012). This approach complements classical tests focusing on short-term observations of a few minutes and/or providing a simple yes/no answer for social preference (Nadler et al., 2004). We documented the magnitude and nature of the social impairments highlighted in the mouse model deleted for the homologous 16p11.2 region generated over a hybrid F1 B6NC3B genetic background (hereafter Del/+). Given the previous study in our laboratory on the same model (Arbogast et al., 2016), we expect Del/+ mice to spent shorter time in contact with others, to follow others less frequently, to approach less and to escape more often the others as well as to emit less ultrasonic vocalisations compared to their wild-type littermates. In addition, as the level of social interactions is related to the activity level, we simultaneously monitor activity and exploration. As most models displayed increased activity over long-term recordings (see supplementary table), we expect Del/+ mice to be hyperactive and display more vertical exploratory behaviour compared to their wild-type littermates. We tested these hypotheses in two contexts of free interactions: interactions between four familiar individuals including one pair of Del/+ mice and one pair of wild-type (wt) mice for males and females separately over three days and three nights (quartet condition) and social encounters of a pair of unfamiliar individuals of the same-genotype over two days and two nights for females only (pair condition; males were not tested in this condition given the higher probability of aggressive behaviour).

## Results

### Quantitative variations in activity and social behaviours

#### Increased locomotor activity but typical vertical exploration

We validated our hypothesis of an increased activity during the active periods of monitoring (i.e., the nights) in both sexes and in both conditions. Indeed, Del/+ animals travelled significantly longer distance compared to their wild-type littermates when recorded in quartets for both males (Linear Mixed Model (LMM) with genotype as a fixed factor and group as a random factor; log-likelihood=-157.3, β=-245.2, SE=89.6, p=0.006) and females (LMM, log-likelihood=-232.2, β=-376.6, SE=155.9, p=0.016; **Figure 1A**) as well as in pairs for females (LMM with genotype as a fixed factor and pair as a random factor, log-likelihood=-262.6, β=-828.5, SE=251.5, p=0.001; **Figure 1B**). In contrast, we did not detect increased vertical exploration in Del/+ mice as suggested by previous studies conducted in new environments with single isolated individuals (Arbogast et al., 2016). Indeed, there was no significant effect of genotype in the number of rearing events when recorded in quartets for both males and females (**Supplementary** Figure 1A) and in pairs for females (**Supplementary** Figure 1B).

**Figure 1:**
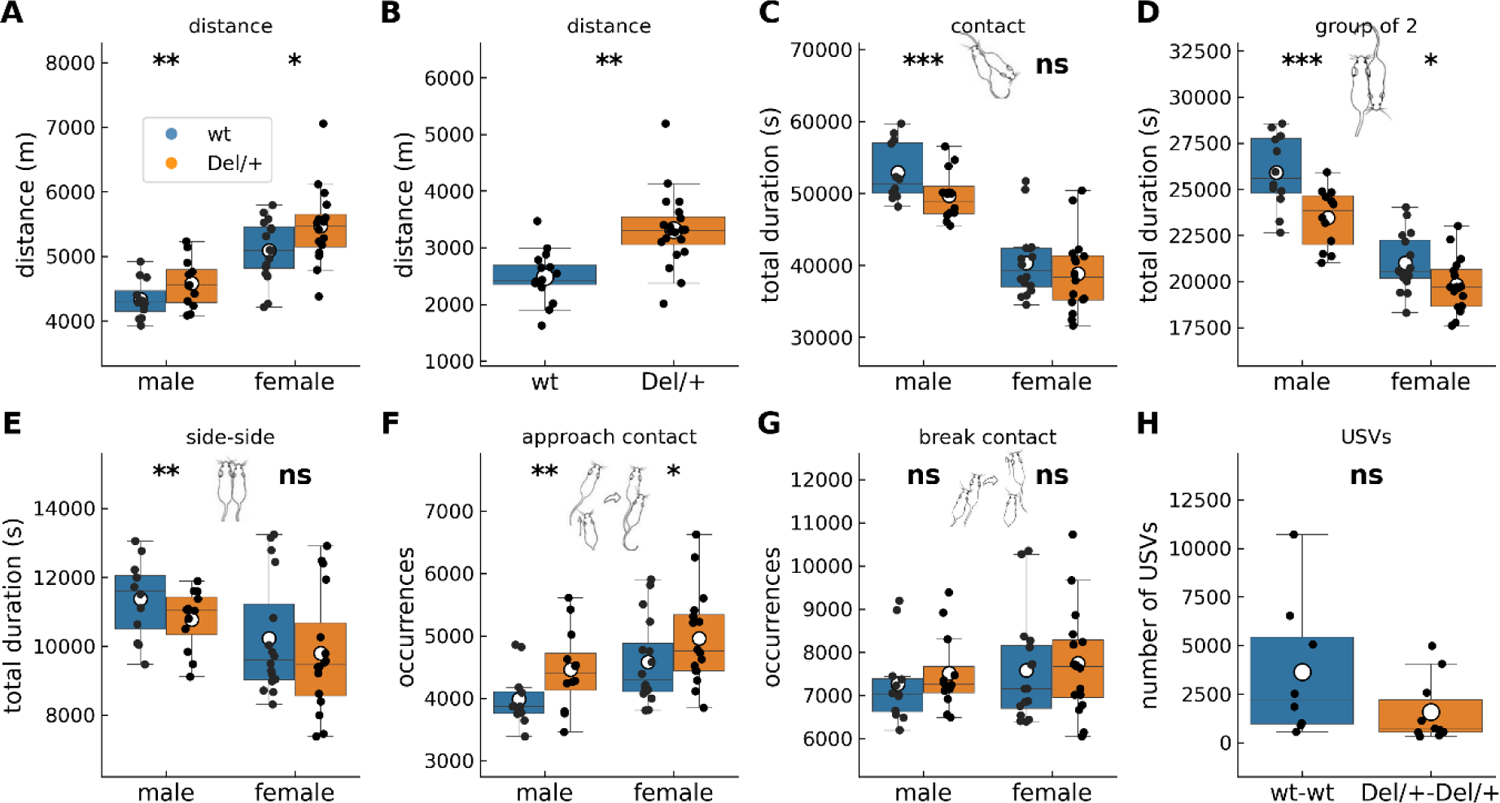
Activity and social behaviours displayed by mice of both sexes and genotypes in long-term monitoring. (A) Distance travelled over the 3 nights of recording of spontaneous behaviours of mixed-genotype quartets of familiar males (12 wt and 12 Del/+ distributed in 6 cages) and females (16 wt and 16 Del/+ distributed in 8 cages). (B) Distance travelled over the 2 nights of recording of spontaneous behaviours of unfamiliar pairs of females of the same genotype (16 wt females distributed in 8 pairs and 20 Del/+ females distributed in 10 pairs). Total time spent in (C) contact, (D) group of 2 mice, and (E) side-side contacts over the 3 nights of recording of spontaneous behaviours of mixed-genotype quartets of familiar males (12 wt and 12 Del/+ distributed in 6 cages) and females (16 wt and 16 Del/+ distributed in 8 cages). Total number of (F) contact initiations and (G) contact terminations over the 3 nights of recording of spontaneous behaviours of mixed-genotype quartets of familiar males (12 wt and 12 Del/+ distributed in 6 cages) and females (16 wt and 16 Del/+ distributed in 8 cages). (H) Total number of ultrasonic vocalisations (USVs) recorded over the 2 nights of recording of spontaneous behaviours of unfamiliar pairs of females of the same genotype (8 pairs of wt females and 10 pairs of Del/+ females). (A-G) Linear Mixed Model, with genotype as fixed factor and group/pair as a random factor; (H) Mann-Whitney U-test; ns: no significant effect of genotype, *: p<0.05, **: p<0.01, ***: p<0.001.

#### Decreased social contacts

When recorded in quartets over three nights, Del/+ males spent shorter time in contact with others (LMM, log-likelihood=-207.3, β=3233.6, SE=821.0, p<0.001; Figure 1C), in contact with one and only one animal (LMM, log-likelihood=-193.5, β=2467.5, SE=460.4, p<0.001; Figure 1D) and in side-by-side contacts (LMM, log-likelihood=-176.2, β=602.7, SE=192.4, p=0.002; Figure 1E) in comparison with their wild-type littermates. Such was not the case of Del/+ females, that displayed only a reduction of time spent in contact with one and only one individual compared to their wild-type littermates (LMM, log-likelihood=-265.6, β=1193.5, SE=524.3, p=0.023; Figure 1D). When recorded in pairs over two nights, unfamiliar Del/+ females did not spend significantly shorter time in contact compared to their wild-type littermates (**Supplementary** Figure 1C). Altogether, we confirmed our hypothesis of reduced social contacts mostly in males; social deficits were subtler in females when observing these global characteristics.

#### Typical follow behaviours

Follow events are rare but well recognisable behaviours that occur mostly when animals are aroused during intense social interactions. These behaviours are therefore more likely to occur in the initial exploration of an unfamiliar conspecific, but they can still be observed between familiar animals housed together (de Chaumont et al., 2021). The number of follow behaviours did not vary significantly between Del/+ and wild-type mice neither in quartets for both sexes (**Supplementary** Figure 1D) nor in pairs for females (**Supplementary** Figure 1E). Therefore, we did not confirm our hypothesis of reduced follow behaviours.

#### Typical approaches and escapes

Comparing approaches and escape behaviours between genotypes is meaningful only when both genotypes are interacting within the same group, otherwise the cage effect could blur the analysis. Therefore, we considered here only quartets. Del/+ mice of both sexes initiated more contacts than their wild-type littermates (Figure 1F), but this was not the case for terminating the contacts (Figure 1G). Therefore, we did not verify our hypothesis of reduced initiation and increased break of contacts.

#### Typical quantity of ultrasonic vocalisations emitted

Ultrasonic vocalisations (USVs) are communicative signals emitted by pups during development and by juvenile and adult mice during social or sexual encounters (Portfors, 2007). At the juvenile or adult stages, USVs reflect the arousal status of the animal and the emotional perception of the interactions (Granon et al., 2018; de Chaumont et al., 2021). As we expected these to be perturbed in Del/+ mice, we hypothesized significant perturbations in the vocal communication of Del/+ pairs compared to wt pairs. Nevertheless, Del/+ pairs of unfamiliar females emitted a similar amount of USVs compared to wt pairs of unfamiliar females (W=60.0, p=0.083; Figure 1H).

Altogether, our initial approach to test hypotheses based on previous studies confirmed perturbations in the activity but only minimal changes in the social and communicative domains. We explored further the social behaviour to detect more subtle abnormalities that could nevertheless deeply impair the everyday-life of our model.

### Qualitative perturbations of social behaviours in quartets

#### Atypical organisation of social behaviours in Del/+ mice

To grab a synthetic view on the behavioural phenotype of the model in quartet condition, we used the centred and reduced data per cage for each Del/+ individual compared to the mean of the whole cage and related to its standard deviation. This Z-score-like measure allows to control the inter-cage variability that we observed in such long-term group recording sessions, while still presenting the whole behavioural profile at a glance. We confirmed the increased activity in both sexes, and the reduced social motivation of Del/+ mice for interactions that was more obvious in males than in females (Figure 2; raw data in **Supplementary** Figures 2-4). Social deficits were highlighted in quartets of both sexes (shorter total time spent in contact, group of 2 and side-side head-to-tail contacts; Figure 2A **& B**). Del/+ male mice also performed less nose-nose, nose-anogenital and side-side (same direction and head-to-tail) contacts compared to their wild-type littermates (Figure 2C **& D**). The structure of the behaviours was affected as the mean durations of contacts and group of 2 events were reduced in Del/+ mice compared to wild-type littermates (Figure 2E **& F**).

**Figure 2:**
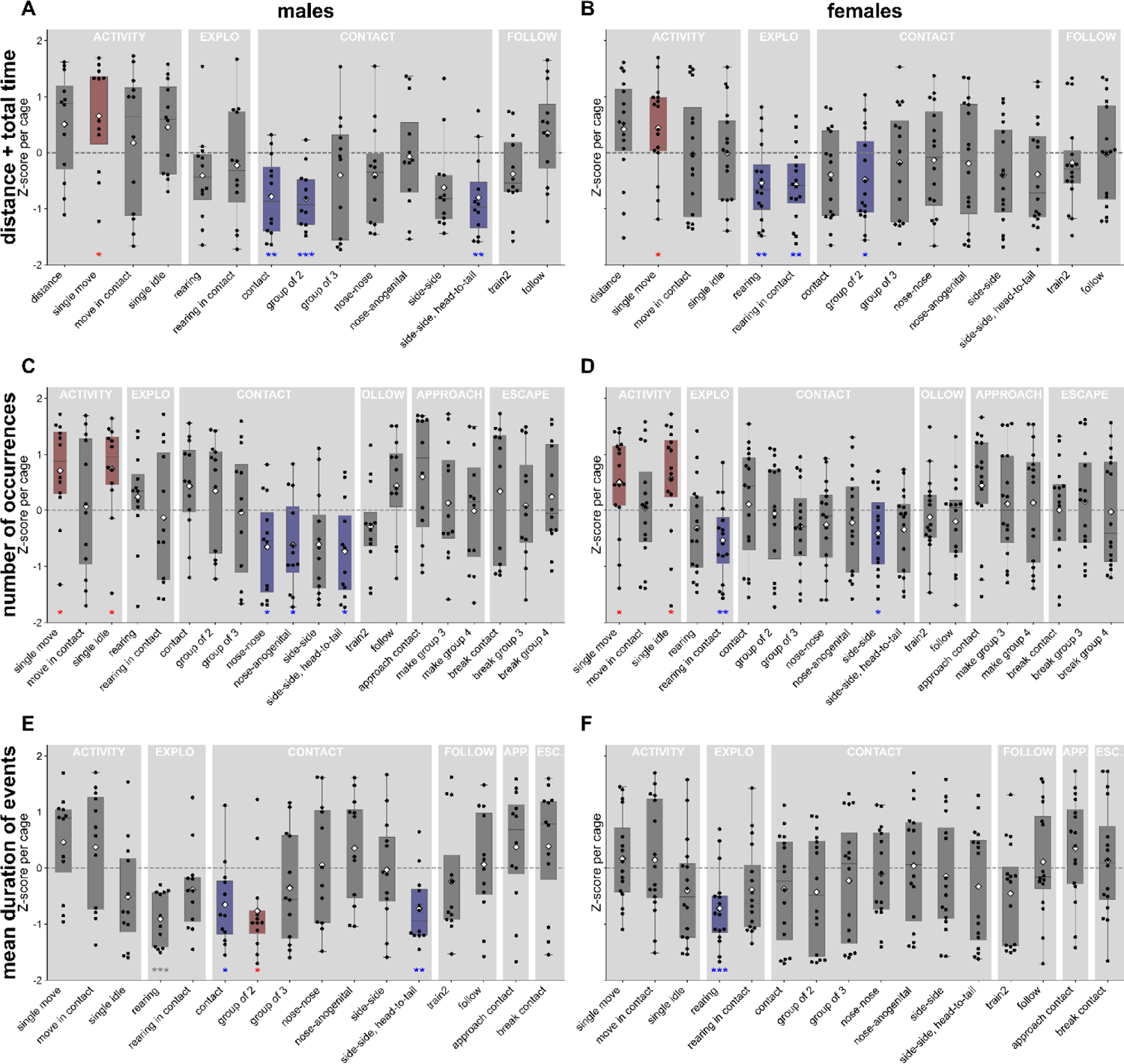
Behavioural profiles for the total duration, the number of occurrences and the mean duration of events of Del/+ males and females over three nights in quartets. (A, B) Z-score profile of the distance and the total duration of each behaviour for each Del/+ mouse compared to the mean behaviour of the four individuals within each quartet for males (A, n=12) as well as for females (B, n=16) over three nights. (C, D) Z-score profile of the number of occurrences of each behaviour for each Del/+ mouse compared to the mean behaviour of the four individuals within each quartet for males (C, n=12) as well as for females (D, n=16) over three nights. (E, F) Z-score profile of the mean duration of behavioural events for each Del/+ mouse compared to the mean value of the four individuals within each quartet for males (C, n=12) as well as for females (D, n=16) over three nights. One-sample t-test if the data follow a normal distribution or Wilcoxon test if data are not normally distributed: *: p<0.05, **: p<0.01, ***: p<0.001. Red boxes figure behavioural events that are significantly more expressed in Del/+ mice compared to the mean of the whole cage; blue boxes depict behavioural events that are significantly less expressed in Del/+ mice compared to the mean of the whole cage; grey boxes reflect non-significant differences between Del/+ and the other animals of the cage.

These structural modifications of contacts displayed by Del/+ mice (especially males) suggest a profound social deficit that could impair the everyday life of the animals, and not just complicate initial encounters with unfamiliar individuals. Indeed, the structural impairments of contacts affect differently the various types of contacts and therefore might impair social maintenance.

#### Selective interactions with wt mice in mixed-genotypes quartets

Quartets involved a pair of wt and a pair of Del/+ mice. Previously, it has been shown in classical laboratory conditions that single-genotype housing of Del/+ mice did not worsen their social phenotype (Yang et al., 2015a). In mixed-genotype conditions, whether mice of one genotype interacted preferentially with mice of the same genotype remains unknown. As wt mice are potentially more socially receptive than Del/+ mice, we expect that individuals of any genotype interact preferentially with wild-type conspecifics. To test this hypothesis, we dissected social interactions between individuals and compared them between mice of the same genotype and mice of the other genotype. Given the group composition (2 wt and 2 Del/+), a mouse of a given genotype has 1/3 of chances to interact with a mouse of the same genotype and 2/3 of chances to interact with a mouse of a different genotype. Therefore, we compared the proportion of time or of occurrences of events with an individual with the same genotype to 1/3. The mean duration of events was compared directly. Over the three nights, both wt and Del/+ males spent a larger proportion of time in contact with a wt individual than with a Del/+ one (t-tests; wt: T=5.07, p=0.0004; Del/+: T=-2.84, p=0.016; Figure 3A), similarly to Del/+ females (t-test; T=-3.12, p=0.007; Figure 3D). wt males and wt females also performed significantly more approaches leading to a contact towards wt mice than towards Del/+ mice (t-tests; males: T=2.35, p=0.038; Figure 3B; females: T=3.51, p=0.003; Figure 3E). In addition, the contacts established with another individual by wt males and females were also significantly shorter when involving a Del/+ mouse than a wt one (Wilcoxon test; males: W=3, p=0.002; Figure 3C; females: W=3, p<0.001; Figure 3F). Overall, the characteristics of social interactions depended on the genotypes involved, with wt mice being more easily approached and maintained in contact compared to Del/+ mice.

**Figure 3:**
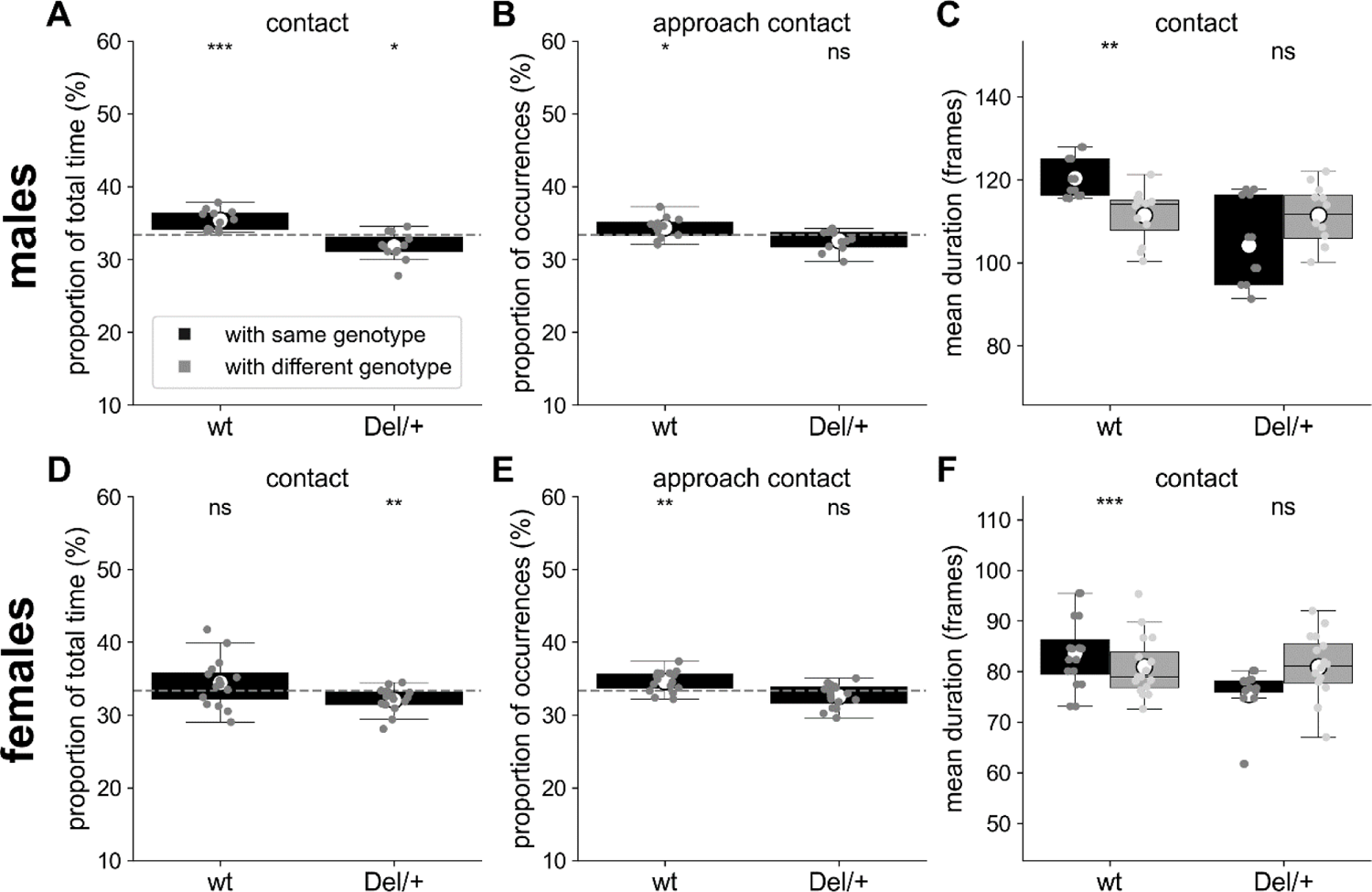
Selective interactions between genotypes for quartets of males and females recorded over three nights. (A) Proportion of the total time spent in contact with individuals of the same or of the different genotype for wt (n=12) and Del/+ (n=12) males. (B) Proportion of the number of approaches leading to a contact with individuals of the same or of the different genotype for wt (n=12) and Del/+ (n=12) males. (C) Mean duration (in frames) of the contacts established with individuals of the same or of the different genotype for wt (n=12) and Del/+ (n=12) males. (D) Proportion of the total time spent in contact with individuals of the same or of the different genotype for wt (n=16) and Del/+ (n=16) females. (E) Proportion of the number of approaches leading to a contact with individuals of the same or of the different genotype for wt (n=16) and Del/+ (n=16) females. (F) Mean duration (in frames) of the contacts established with individuals of the same or of the different genotype for wt (n=16) and Del/+ (n=16) females. (A, B, D, E) One-sample t-tests compared to expected proportions; dashed horizontal lines represent the expected proportions: 1/3 with individuals of the same genotype. (C, F) Non-parametric Wilcoxon paired tests. ns: not significant, *: p<0.05, **: p<0.01, ***: p<0.001.

### Qualitative perturbations of social behaviours in pairs

#### Atypical organisation of social behaviours in pairs of Del/+ mice

In pairs of females, over the two nights of recording, Del/+ female pairs established significantly more contacts per hour (MW: U=15, p=0.027; **Supplementary** Figure 5) of shorter mean duration (MW: U=70, p=0.006; **Supplementary** Figure 5) compared to their wt littermates. This perturbed temporal structure of global contact might be related to the increased activity of the Del/+ mice compared to wt mice. Nevertheless, this increased activity compared to wt mice affected mostly other contacts than the specific ones (nose-nose, side-side, and side-side head-to-tail), that did not differ between genotypes.

Del/+ females performed follow behaviours at a lower speed and travelled less distances during these behaviours compared to wt females (see **Supplementary** Figure 6 for train2; such an effect was not significant in follow behaviours without ano-genital contacts (data not shown)). These qualitative variations suggest that Del/+ mice displayed decreased social motivation and arousal compared to wt mice over long-term experiments.

## Shortened sequences and atypical acoustic structure of ultrasonic vocalisations in Del/+ mice

We next explored the qualitative variations of ultrasonic communication between Del/+ pairs and wt pairs. USVs are organised in sequences, i.e., consecutive USVs separated by less than 750 ms (de Chaumont et al., 2021). These sequences were significantly shorter in Del/+ pairs compared to wt pairs (LMM: n(wt)=2923, n(Del/+)=2578, β=4.378, SE=1.242, p=0.002; Figure 4A). This might reflect the reduced arousal of Del/+ pairs compared to wt pairs in the maintenance of these interactions. When testing the effect of the deletion on variables describing the acoustic structure (Figure 4B), we observed that USVs recorded over the two nights from Del/+ pairs (n=15885) were acoustically simpler, with shorter duration (Figure 4C), a smaller frequency range covered (Figure 4D), less frequency modulations (Figure 4E) compared to those recorded in wt pairs (n=29079); frequency characteristics (Figure 4F) and frequency jumps (data not shown) did not differ between genotypes.

**Figure 4:**
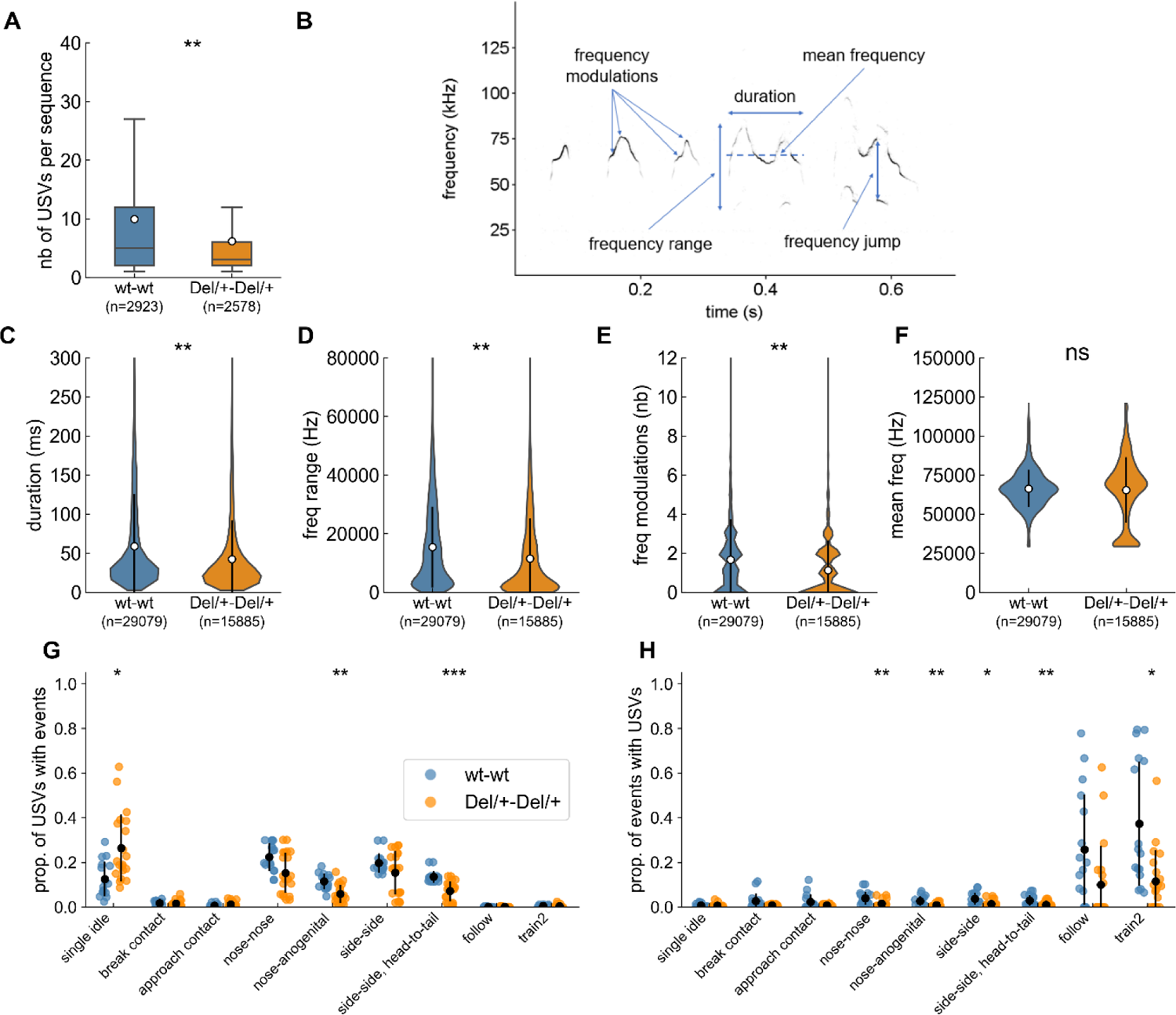
Characteristics of ultrasonic vocalisations (USVs) emitted during social encounters between two unfamiliar females of the same genotype. (A) Number of USVs per sequence (i.e., successive USVs separated by less than 750 ms) estimated over 2 nights; sample sizes represent the number of sequences analysed and the white dots represents the mean values. (B) Spectrogram (300 kHz sampling frequency, 16-bits format, FFT length: 1024 points, 75% overlap, Hamming window, 0.853 ms time resolution) of five USVs depicting the main acoustic features measured by LMT USV Toolbox. (C) Duration of USVs. (D) Range of frequencies covered by USVs. (E) Number of frequency modulations (i.e., inflexion points in the peak frequency) per USV. (F) Mean peak frequency computed over each USV. (C-F) sample sizes represent the number of USVs analysed; the white dot and black error bars represent the mean values and standard deviations. (G) Proportion of the total number of USVs recorded occurring synchronously with the different behavioural events over the two nights. (H) Proportion of events occurring synchronously with USVs over the two nights. (A-F) linear mixed model with genotype as fixed factor and pair as random factor; (G-H) Mann-Whitney U-tests between 8 wt-wt pairs and 10 Del/+-Del/+ pairs. ns: not significant, *: p<0.05, **: p<0.01, ***: p<0.001.

The call rates specific to behavioural events did not differ significantly between wt and Del/+ pairs, with the lowest call rate in single idle and the highest in the different types of contacts, (**Supplementary** Figure 7). Del/+ pairs emitted significantly more USVs in the single idle context and significantly less USVs in nose-anogenital contact, and in side-side head-to-tail contacts compared to wt pairs (Figure 4G). This reflects the reduced expression of these behaviours in Del/+ mice compared to wt mice. The proportion of events that were accompanied by USVs was significantly reduced in Del/+ pairs compared to wt pairs for nose-nose contacts, nose-anogenital contacts, side-side contacts, side-side head-to-tail contacts and train2 events (Figure 4H). Altogether, this suggests that Del/+ mice were less aroused by intense social contacts.

### Atypical sequences of social contacts in Del/+ pairs

As the structure of behavioural events such as contacts was perturbed in Del/+ mice (see above), we investigated whether the organisation of the behaviour was affected by examining the temporal succession of simple behavioural events, that should contribute to a better understanding of the functions of the different behaviours (Wiltschko et al., 2015; Bels et al., 2022). For this analysis, we defined exclusive behavioural events (i.e., events that could not occur at the same time; see methods for definitions). We focused on simple behavioural blocks to explore the bases of the behaviour: the animal is alone and moving or idling, and the different types of contacts; more complex social events combining specific movements and specific types of contact or proximity such as follow or train2 were not examined in the present analysis. The exclusive behavioural events were computed by separating the existing non-exclusive events. We excluded any overlap between events and each animal of the pair was engaged in one and only one event at each time frame.

Recomputing the behavioural profiles with these exclusive events allowed to specify social contact deficits: Del/+ pairs displayed structural variations (mean duration) of events involving side-side contacts (and only side-side contacts) between genotypes (**Supplementary** Figure 8). To analyse the temporal succession of events, we compared the transitions from one behavioural event to another between pairs of wt females (Figure 5A) and pairs of Del/+ females (Figure 5B).

**Figure 5:**
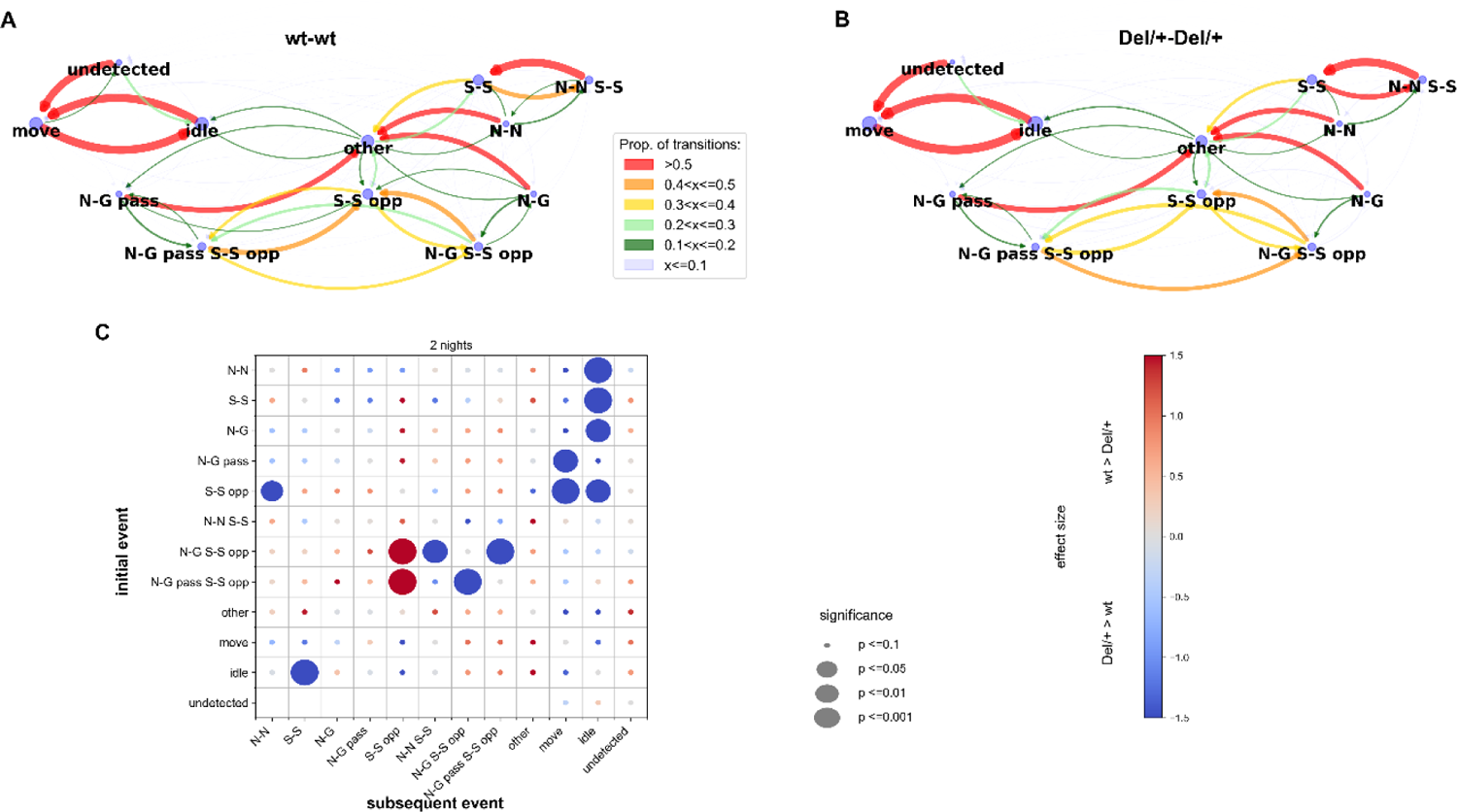
Transitions between exclusive behavioural events in unfamiliar female pairs. (A,B). Examples of the transitions between behavioural events in pairs of wt (A) and in pairs of Del/+ (B) females over the long-term recording. The proportion of each possible transition is represented by the colour and weight of the arrows oriented from initial to subsequent events. The size of the blue nodes represents the proportion of each event. (C) Overview of the comparisons between wt-wt and Del/+ Del/+ pairs of the transitions for the events in the y-axis towards the events of the x-axis over the two nights of recordings. n(wt-wt)=16, n(Del/+ Del/+)=20; Mann-Whitney U-tests; significance levels are represented by the diameter of the circles, and the effect size is represented by the colour of the points. N-N: nose-nose contact, N-G: nose-anogenital contact, N-G pass: passive nose-anogenital contact, S-S: side-side contact, S-S opp: side-side contact head-to-tail, N-N S-S: nose-nose contact during side-side contact, N-G S-S opp: nose-anogenital contact during side-side contact head-to-tail, N-G pass S-S opp: passive nose-anogenital contact during side-side contact head-to-tail, other cct: other types of contacts than the ones described above, idle: single idle, move: single move, undetected: the animal is not detected.

Over the two nights of recording, Del/+ mice appeared to show more transitions back and forth between ‘side-side head-to-tail & ano-genital sniffing’ (being sniffed or sniffing; N-G S-S opp or N-G pass S-S opp), more transitions between ‘side-side head-to-tail & ano-genital sniffing’ (N-G S-S opp) and ‘nose-nose & side-side’ (N-N S-S), and less transitions between ‘side-side head-to-tail & anogenital sniffing’ (N-G S-S opp) and pure ‘side-side head-to-tail’ (S-S opp) compared to wt mice, as if Del/+ mice performed more continuous ano-genital sniffing during side-side head-to-tail behaviours. Del/+ mice displayed an atypical start of a social sequence: they used side-side contacts (S-S) as a social sequence start (i.e., following an idle event) more frequently compared to wt mice. In addition, social sequences appeared to end in an atypical way in Del/+ mice since nose-nose (N-N), nose-anogenital (N-G), passive ano-genital (N-G pass), side-side (S-S), and side-side head-to-tail (S-S opp) ended a social sequence (i.e., were followed by idle or move) more frequently in Del/+ mice compared to wt mice (Figure 5C). Altogether, this suggests that the perturbations of the behavioural sequence in Del/+ mice concerned the initiation and termination of social contacts but did not affect the most frequent transitions between the different types of contacts.

## Discussion

In the present study, Del/+ mice displayed differential impairments according to sex over long-term monitoring (Figure 6). The expected hyperactive phenotype appeared in both sexes. In contrast, in the social domain, among familiar quartets, Del/+ males displayed reduced social interactions, while these deficits were subtler in Del/+ females. Interestingly, the behavioural variations were perceived by the animals themselves, as both wt and Del/+ mice displayed a social preference toward wt animals. In encounters between unfamiliar females, Del/+ mice showed mostly qualitative variations in ultrasonic vocalisations and in the organisation of their social interactions compared to wild-type mice.

**Figure 6:**
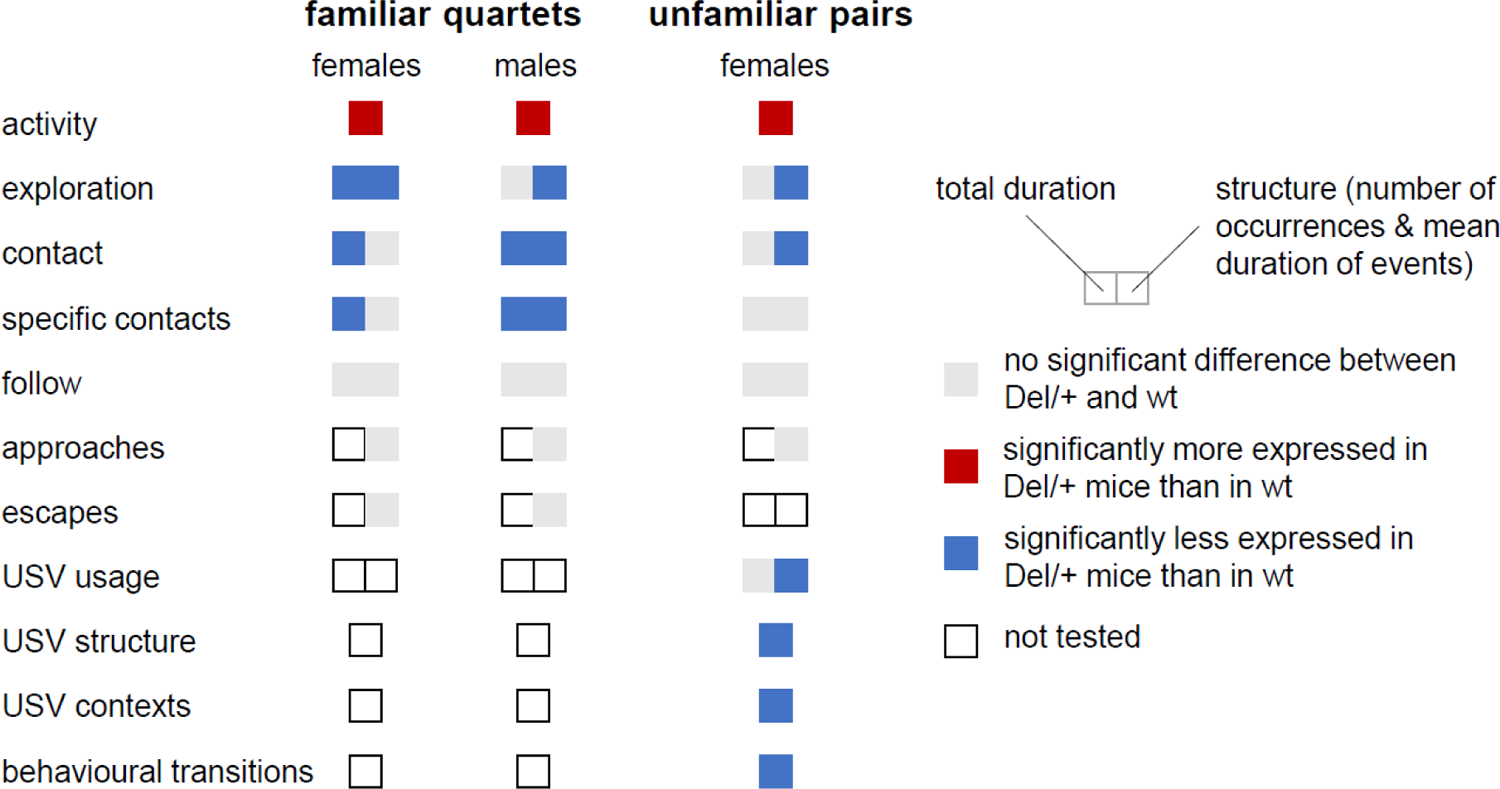
Summary of behavioural variations between Del/+ and wild-type mice. Variations were explored in both sexes and in both contexts (familiar quartets and unfamiliar pairs). Variations between genotypes are depicted in nuances of grey as detailed in the legend. The left part of the coloured rectangles represents variations in total duration and the right part represents variations in the structure of the events (e.g., number of events and in mean duration).

### Disentangling activity and social phenotypes

As activity and exploration might be traits which can affect social behaviours, we tested them simultaneously with social behaviours. Our comparison of genotype-related differences in activity in both sexes confirmed previous findings on the same model in different protocols.

Indeed, in the study of Arbogast using the same model, the hyperactivity displayed by Del/+ mice was visible only over the dark phase in the circadian activity test, while it was not observed in the 30 min exploration of an open field (Arbogast et al., 2016). In contrast to our previous study in which we observed increased rearing over long-term in isolated Del/+ individuals (Arbogast et al., 2016), we observed a different pattern for vertical exploration, with a decreased number of rearing events in Del/+ mice, in quartets of both sexes and in pairs of females. Reduced muscle strength can be ruled out (Arbogast et al., 2016). This discrepancy might be related to differences in the test cage: in the present study, mice were tested in a social context with bedding and nesting material, while in (Arbogast et al., 2016) mice were isolated in a new environment without bedding and nesting material. Reduced unsupported rearing is expected in more anxious animals (Sturman et al., 2018). However, as the proportion between supported and unsupported rearing did not vary significantly between genotypes (data not shown), other causes than increased stress and anxiety in Del/+ mice of the present study remain to be investigated. The increased activity displayed by Del/+ mice might explain the shorter mean duration of rearing events over long-term monitoring. Over the long-term, impairments in activity and exploration occurred simultaneously with social deficits in our testing conditions. In this case, we cannot rule out an influence of hyperactivity on social deficits, as in some other mouse models of autism (e.g., *Shank2/ProSAP1^-/-^ mice*: (Schmeisser et al., 2012; de Chaumont et al., 2019)).

### Decoding social defects in unfamiliar pairs

We observed that the deletion impairs the social life in our mice. In initially unfamiliar pairs, when the animals get familiar with each other over the long term, mutant mice displayed qualitative impairments such as atypical ways of starting and ending contact sequences. In more classical phenotyping studies, short-term interactions are monitored and these provide different phenotypes. These changes in phenotypes might follow the evolution of the behaviours of mice over time (visible burrow system, (Arakawa et al., 2007)). Indeed, in our case, in the first 15 min of interaction of the pairs of unfamiliar females, we observed simple quantitative reduction of time spent in contact and follow behaviours in Del/+ mice compared to wt mice (**Supplementary** Figures 8 **& 9**) when the animals are still unfamiliar to each other. Observations over the short-term parallel classical tests for social interactions, with a decreased time spent sniffing the conspecific (males; (Arbogast et al., 2016)) and reflect the atypical way of initiating social encounters, in which ano-genital sniffing and following appeared to play a crucial role. In contrast, observations of structural abnormalities in the social behaviour over the long-term reflect the difficulties in maintaining social interactions (Chevallier et al., 2012), which has been under-studied up-to-now given the short duration of classical social experiments. Such structural defects might be more complicated to improve through behavioural intervention (as in (Pujol et al., 2018)) compared to motivational defects and require further study for a better understanding of the neuronal circuits involved. Future studies will unravel the time course of social interactions to identify the time point at which the initial social contacts turn to social maintenance.

### Sex differences in the social phenotype

These social investigations could not be run fully in both sexes since we were not able to run the social encounters between unfamiliar males. Indeed, sexually mature males are highly aggressive and could not be left for two days and two nights together without severe fighting outcomes despite the large surface of the test cage (personal observation). We nevertheless observed robust social impairments in males over the long-term when tested with familiar cage mates. The reductions of duration, number and mean duration of some specific contacts were even stronger in males than in females, while the activity level was increased in Del/+ mice of both sexes to a similar extent. Interestingly, the fact that social deficits were already visible over the short-term in females recorded in unfamiliar pairs (**Supplementary** Figures 8 **& 9**) might reflect the fact that Del/+ females might be vulnerable to the combined stress of social unfamiliarity and of the new physical environment as it has been found in another model (Giovanniello et al., 2021). These findings might be reminiscent of observations in patients. Indeed, in patients carrying a 16p11.2 deletion, the sex ratio was almost balanced, with 1.3 males for 1 female for autism and 1.6 male for 1 female for ID/DD.

However, females carrying a 16p11.2 deletion displayed comorbid features more frequently than males (Polyak et al., 2015). There was an increased tendency of female patients to display anxiety-like disorders (discussed in (Giovanniello et al., 2021), which might also affect the diagnosis of patients (Dean et al., 2016; Beggiato et al., 2017)). Future studies should improve the testing conditions to be able to test also males in this condition of combined stress of social and environmental unfamiliarity.

### Effect of familiarity with the environment

This susceptibility to anxiety in females might also be brought closer to the reaction to the environment. As suggested by previous studies (Portmann et al., 2014), the Del/+ mice might have difficulties in habituating to new environments. Robust social deficits were observed over the long term; at this time scale, we can suppose that wild-type mice got habituated while Del/+ mice did not. Over the short-term (first 15 min), Del/+ mice of both sexes did not show social deficits in our conditions of familiar cage mates (**Supplementary** Figure 10). In this case, the arousal triggered by the environmental change might mask social impairments over the short-term experiment. In contrast, when interacting with an unfamiliar conspecific, Del/+ females displayed quantitative reduction of social contacts over the short-term already (**Supplementary** Figure 9). This suggests that initial social encounters might be even more stressful for Del/+ female mice (see above). This increased behavioural reaction might also be triggered by the fact that these mice were tested in unfamiliar pairs in the testing environment that they already visited for the recordings in quartets. Such a re-exposure to the unfamiliar testing environment might boost behavioural deficits, as in the *Shank3^ΔC/ΔC^*mice (Krüttner et al., 2022). To explore this aversion towards unfamiliarity, further studies should incorporate social cognitive challenges in long-term monitoring of mixed-genotype groups within a complex changing environment to better fit the natural needs of mice (Gray et al., 2000) and to provide cognitive tasks to unravel social phenotypes (Winiarski et al., 2021, 2022).

### Atypical ultrasonic vocalisations

Mouse ultrasonic vocalisations cannot be considered as direct proxies for speech abnormalities since they are mostly innate (Mooney, 2020). In 16p11.2 Del/+ mouse models, vocal production impairments were minimal. Indeed, previous studies highlighted that Del/+ mice were able to utter all types of ultrasonic vocalisations in adults (Portmann’s model: (Yang et al., 2015b)) and in pups (Horev’s model: (Agarwalla et al., 2020)). In our study, we only observed slight variations in the acoustic structure of the USVs, which might reflect a simplification of the calls (shorter, less frequency modulated). The reduction of usage that we observed (less USVs, in shorter sequences) might reflect more closely the reduced arousal during social interactions in Del/+ mice compared to wild-type mice. This corroborates and refines behavioural findings and represents a proxy for social arousal (de Chaumont et al., 2021), probably more than articulation or phonological errors or dysarthria identified in most patients carrying a 16p11.2 deletion (Rosenfeld et al., 2010; Mei et al., 2018).

## Perspectives

The characterisation of the present model highlights robust social deficits, that also seems to parallel sex-related variations in patients. The same framework could be used to examine the contribution to the social phenotype of specific brain regions, with a focus on the striatum as a key hub structure associated with action selection, cognitive flexibility, attention, sensory selection, reward processing and goal-directed behaviours (Fuccillo, 2016), all these functions being involved in social interactions. Indeed, as the dorsal medial striatum is associated with goal-directed behaviours, the dorsal lateral striatum with automated behaviours and the nucleus accumbens with motivational states and reward processing, the 16p11.2 deletion in each of these striatal regions will document their involvement in the different social deficits observed. In this same line, the contribution to the phenotype of each gene within the deleted region could also be detailed, as it has been done to identify the contribution of *Kctd13* gene to the cognitive impairment phenotype (Arbogast et al., 2019; Martin Lorenzo et al., 2021). To ascertain the robustness of these findings, a cross-species comparison should be conducted in the rat model (Martin Lorenzo et al., 2023), as recommended in recent guidelines to increase the value and robustness of preclinical models (Silverman et al., 2022). Rescue strategies could then be attempted, with for instance R-baclofen, a GABAb agonist, or Fasudil, an inhibitor of the Rho-associated protein kinase, both restoring the cognitive deficits in the mouse model (Stoppel et al., 2018; Martin Lorenzo et al., 2021). Currently, the effects of such treatment on the social phenotype is not documented and it would be of interest to evaluate its therapeutic value.

## Material and methods

### Animals

Mice were generated according to the breeding scheme used in (Arbogast et al., 2016). In brief, C57BL/6N.16p11.2 Del/+ females were bred with sighted C3H/HeH (C3H.*Pde6b^+^* noted here C3B) males (Hoelter et al., 2008) (16p11.2+/+) to obtain F1 C57BL/6N x C3B.16p11.2 Del/+ (hereafter Del/+) and F1 C57BL/6N x C3B.16p11.2 +/+ (hereafter wt) mice. The cohort included 24 males (12 wt and 12 Del/+) and 32 females (16 wt and 16 Del/+). Animals were grouped in cages of four animals at weaning (quartets: 2 wt and 2 Del/+), therefore leading to 6 cages of males and 8 cages of females. In addition, for paired social encounters, we added two pairs of Del/+ females of the same age and housed in similar conditions. All mice were housed under 21-23°C with 12h/12h light/dark cycle (lights on at 7:00 AM). Hemp squares, food and water were available ad libitum. All mice were weighted at 11 weeks.

### Individual identification

Mice were identified through finger cuts realised between 2 and 7 post-natal days. Genotyping was conducted on these finger biopsies according to the protocol described in (Arbogast et al., 2016). In brief, DNA was extracted in NaCl. PCR reaction used the primers Del70 F (CCTGTGTGTATTCTCAGCCTCAGGATG) and primer Del71 R (GGACACACAGGAGAGCTATCCAGGTC) with the following cycles: one cycle of 4 min at 95°C, 35 cycles of 30°C at 94°C + 30 s at 62°C + 1 min at 72°C, one cycle of 7 min at 72°C. At least two weeks before starting the recordings, we inserted a Radio Frequency IDentification (RFID) tag (APT12 PIT tags; Biomark, Inc., Boise, The United States of America) under the skin of each individual under gas anaesthesia (Isoflurane) with local analgesia (Lidor 20 mg/ml, with 40 ul/10 g mouse). RFID tags were located in the lower part of the left flank. Mice were allowed to recover for one week. They were manipulated three days before starting the behavioural experiments to get them used to the experimenters and to being held within a cup. Mice were habituated to the experimental room and the setup since they underwent the novel object recognition test (data not presented) in the same room and setup at least one week before the quartet recordings. They underwent the dyadic encounters at least one week after the quartet recordings, and were therefore also familiar with the experimental room.

### Behavioural monitoring in quartets

We monitored the individual and social behaviours of each quartet of mice over three days and nights in the Live Mouse Tracker system (LMT, plugin 931; (de Chaumont et al., 2019)). This system tracks individually mice living in a group over several days and nights and extracts automatically the number, total duration and mean duration of more than thirty behavioural events describing the posture of the mouse, the types of social contacts, the dynamic social approach and escapes as well as complex social groupings (see (de Chaumont et al., 2019)). In this system, the four mice (10-14 weeks of age) of each housing cage were left undisturbed for 71 hours in a large transparent Plexiglas cage (50 x 50 x 40 cm), with fresh bedding, a house (width: 100 mm, depth: 75 mm, height: 40 mm) in red Plexiglas, 6 dental cotton rolls as well as food and water ad libitum. Light/dark cycle and temperature conditions were similar to those of the housing room (12/12h light/dark, lights on at 07:00 AM, 75-90 lux when the lights were on). Each recording session started between 03:00 and 04:00 PM. At the end of the session, mice were placed back in their home cage and the LMT setup was cleaned with soap water and dried with paper towels. Altogether, we recorded the six cages of males and the eight cages of females, keeping the animals with their familiar cage mates. For each individual, we extracted the total distance travelled. We also automatically recorded the following behavioural events (based on the original publication of LMT (de Chaumont et al., 2019); the type of quantification extracted is indicated in brackets):

**Single move**: The focal animal is moving (speed > 5 m/s) without being in contact with any other animal (total duration, number of events, mean duration of events).

**Move in contact**: The focal animal is moving (speed > 5 m/s) while being in contact with another animal (total duration, number of events, mean duration of events).

**Jumps**: The focal animal is jumping against the wall (total duration, number of events, mean duration of events).

**Single idle**: The focal animal is resting (not moving) without being in contact with any other animal (total duration, number of events, mean duration of events).

**Rearing**: The focal animal is straightened on its hindlegs (either unsupported or against the wall). Rearing is considered when the body slope is higher than a threshold (total duration, number of events, mean duration of events).

**Rearing in contact**: The focal animal is straightened on its hindlegs (either unsupported or against the wall) while being in contact with another individual. Rearing is considered when the body slope is higher than a threshold (total duration, number of events, mean duration of events).

**Contact**: The focal animal is touching another individual (total duration, number of events, mean duration of events).

**Group of 2**: The focal animal is touching one and only one other individual (total duration, number of events, mean duration of events).

**Group of 3**: The focal animal is touching two and only two other individuals (total duration, number of events, mean duration of events).

**Nose-nose**: The focal animal is sniffing the nose of another animal (i.e., the nose is at a whisker distance from the nose of the other animal) (total duration, number of events, mean duration of events).

**Nose-anogenital**: The focal animal is sniffing the ano-genital region of another animal (i.e., the nose is at a whisker distance from the tail basis of the other animal) (total duration, number of events, mean duration of events).

**Side-side**: The flank of the focal animal is in contact with the flank of another animal; both animals head in the same direction (total duration, number of events, mean duration of events).

**Side-side head-to-tail**: The flank of the focal animal is in contact with the flank of another animal; both animals head in opposite directions (total duration, number of events, mean duration of events).

**Train2**: The focal animal is moving (speed > 5 m/s) while sniffing the ano-genital region of another animal also moving (total duration, number of events, mean duration of events).

**Follow**: The focal animal is walking in the path of another individual: the two animals are moving at a speed >5 cm/s, the angles between the two animals are less than 45° apart, and the mass centre of the follower (the focal animal) is within a follow zone of one mean body length of width and two mean body lengths of length (total duration, number of events, mean duration of events).

**Approach contact**: The focal animal gets closer to another one, with the approaching animal walking at a higher speed than the approached animal; the approach ends by a contact between the two animals (total duration, number of events, mean duration of events).

**Make group3**: The focal animal is joining a group of two animals to form a group of three animals in contact (number of events).

**Make group4**: The focal animal is joining a group of three animals to form a group of four animals in contact (number of events).

**Break contact**: The focal animal is getting away (higher speed) from the animal it has been in contact with; the speed of the focal animal is higher than the speed of the other animal (number of events).

**Break group3**: The focal animal is leaving a group of three animals to leave a group of two animals in contact; the focal animal has the highest speed among the three animals in contact (number of events).

**Break group4**: The focal animal is leaving a group of four animals, that remain as a group of three animals in contact; the focal animal has the highest speed among the four animals in contact (number of events).

For social events, we computed the variables either in general or separately according to the identity of the interacting individual. These behaviours are not exclusive: one animal can be involved in several of them simultaneously.

### Social encounter between unfamiliar individuals in pairs

We evaluated the social interactions and communication between unfamiliar individuals in pairs. For these recordings of social behaviour and ultrasonic communication, we focused on pairs of individuals since we currently cannot identify the emitter of USVs when animals were interacting closely. Therefore, we recorded undisturbed dyadic interactions between two unfamiliar individuals (from two different housing cages) of the same age (14-20 weeks of age) and genotype for 47h (two days and nights, starting between 03:00 and 04:00 PM). For that purpose, we coupled the LMT system (plugin 931) with the Avisoft Ultrasound Gate 416 (300 kHz sampling rate, 16-bit format; trigger: level of this channel; pre-trigger: 1 s; hold time: 1 s; duration > 0.005 s; trigger event: 2 % energy in 25-125 kHz with entropy < 50%; Avisoft Bioacoustics, Glienecke, Germany) connected to a CM16/CMPA microphone (Avisoft Bioacoustics, Glienecke, Germany). LMT and Avisoft systems were synchronised based on the protocol described in (de Chaumont et al., 2021). Altogether, we recorded eight pairs of wt females and ten pairs of Del/+ females. We focused on females since males were too aggressive toward each other when they were taken out of their housing group to conduct robust (and safe) social monitoring. We recorded the same behaviours as in quartets recordings, except those involving more than two animals. USVs were analysed using LMT – USV Toolbox (de Chaumont et al., 2021).

### Transitions between exclusive behavioural events

To investigate the transitions between two events in paired encounters, we needed to compute exclusive events, i.e., events that do not overlap in time for each individual. For that purpose, we split the existing overlapping events in simpler events that were not overlapping in time to obtain new exclusive events (script ComputeTransitionsBetweenEvents.py). We obtained the following exclusive events:

**Move:** The focal animal is moving (speed > 5 m/s) without being in contact with any other animal.

**Idle:** The focal animal is resting (not moving) without being in contact with any other animal.

**Nose-nose**: The focal animal is sniffing the nose of another animal (i.e., the nose is at a whisker distance from the nose of the other animal).

**Nose-anogenital**: The focal animal is sniffing the ano-genital region of another animal (i.e., the nose is at a whisker distance from the tail basis of the other animal).

**Passive nose-anogenital**: The focal animal is being sniffed in the ano-genital region by another animal (i.e., the nose is at a whisker distance from the tail basis of the focal animal).

**Side-side**: The flank of the focal animal is in contact with the flank of another animal; both animals head in the same direction.

**Side-side head-to-tail**: The flank of the focal animal is in contact with the flank of another animal; both animals head in opposite directions.

**Nose-nose & Side-side**: The focal animal is sniffing the nose of the other animal during a side-side contact with this same animal.

**Nose-anogenital & side-side head-to-tail**: The focal animal is sniffing the ano-genital region of the other animal during a side-side head-to-tail contact with this same animal.

**Passive nose-anogenital & side-side head-to-tail**: The focal animal is being sniffed in the ano-genital region by the other animal during a side-side head-to-tail contact with this same animal.

**Other contact**: The focal animal is in contact with another animal and this type of contact is not one of the above described ones (i.e., nose-nose, nose-anogenital, side-side, side-side head-to-tail, nose-nose & side-side, or nose-anogenital & side-side head-to-tail).

**Undetected**: The focal animal is not detected (tracking issues). This event was needed to have each animal engaged in one event at each time frame.

We computed the proportion of transitions ‘A to B’ from one event (event A) to another (event B) by dividing the number of transitions ‘A to B’ by the total number of occurrences of event A. This was conducted for each individual separately, as each individual was involved in one and only one event at each moment.

### Statistical analyses

We did not exclude any outlier. For behaviours in quartet monitoring and at the individual level during paired encounters (e.g., activity, exploration, asymmetric social events), and acoustic features of USVs and USV sequences, we used linear mixed models (LMM; *mixedlm()* function from the *statsmodels 0.13.2* package in Python 3.8), with genotype as fixed factor and cage (either quartet or pair) as random factor. This type of analyses was robust to violation of assumptions for distribution (Schielzeth et al., 2020). LMM results are presented with the log-likelihood as the goodness of fit, β as the coefficient estimate of the fixed factor ‘genotype’, i.e., the slope of the line between wt and Del/+ values, SE as the standard error of this coefficient estimate and p as the probability of the current data to occur assuming the difference between genotypes is null).

For the behavioural profiles computed in quartets, we also conducted an analysis with centred and reduced data of the Del/+ mice per cage (i.e., per quartet) and compared these z-score values for each Del/+ individual to 0 using Student’s one-sample t-tests (*ttest_1samp()* function from the *SciPy 1.8.0* package of Python 3.8) if the data were normally distributed (Shapiro-Wilk test, *shapiro()* function from the *SciPy 1.8.0* package of Python 3.8) and Wilcoxon’s test if not (*wilcoxon()* function from the *SciPy 1.8.0* package of Python 3.8). We chose this option in addition to comparing raw data in order to consider the important variations between quartets in such conditions and to present the whole behavioural profile at a glance for each condition. This presentation is even more stringent than testing row data (that are presented in the first result sections and in **Supplementary** Figures 2-4) given the comparison with the mean of the whole cage and not just with wt animals. The total duration and number of occurrences of selective interactions according to the genotype of the individuals were first tested for normality (Shapiro-Wilk test, *shapiro()* function from the *SciPy 1.8.0* package of Python 3.8) and equality of variances (Fisher-Snedecor test, *f()* function from the *SciPy 1.8.0* package of Python 3.8). As data were normally distributed and had equal variances, we compared data to the chance level to interact with individuals of the same genotype (1/3; see result section) using Student’s T-tests (*ttest_1samp()* function from *Scipy 1.8.0* package of Python 3.8); the mean duration of these selective interactions was compared using T-test when they were normally distributed and had equal variance (*ttest_ind()* function from *Scipy 1.8.0* package of Python 3.8) or paired Wilcoxon tests when this was not the case (*wilcoxon()* function from *Scipy 1.8.0* package of Python 3.8).

Given the small sample sizes of our data for social behaviours at the pair level (e.g., contact, nose-nose contact, side-side contact, side-side head-to-tail, total number of USVs) in encounters between unfamiliar individuals, we used non-parametric Mann-Whitney U tests from the *SciPy 1.8.0* package of Python 3.8. Proportion of transitions between exclusive behavioural events were compared at the individual level between genotypes using Student’s T-test if data were normally distributed (Shapiro-Wilk test) or Mann-Whitney U-tests in other cases (tests from the *SciPy 1.8.0* package of Python (3.8)). In this case, P-values were corrected by the number of tests conducted (12*11) and effect size was estimated on raw data using the Cohen’s D indicator. All scripts are available (https://github.com/fdechaumont/lmt-analysis).

## List of abbreviations

RFID: Radio Frequency Identification

LMT: Live Mouse Tracker

USV: UltraSonic Vocalisation

LMM: Linear Mixed Model

>MW: Mann-Whitney U-test

## Declarations

### Ethics approval

In compliance with ethical rules and regulatory requirements on use and welfare of laboratory animals, this protocol and all procedures have been reviewed and approved by our ethical committee (Com’Eth, CE17) registered at the French Ministry of Research under the reference: 2018062715092398 (DAP15692).

### Consent for publication

Not applicable.

### Availability of data and materials

The datasets generated and/or analysed during the current study are available in the Zenodo repository, [PERSISTENT WEB LINK TO DATASETS here available upon publication]. Ultrasonic vocalisations data will also be available on the MouseTube database (53).

## Funding

This work was supported by the National Centre for Scientific Research (CNRS), the French National Institute of Health and Medical Research (INSERM), the University of Strasbourg (Unistra), and the French government funds through the “Agence Nationale de la Recherche” in the framework of the Investissements d’Avenir program [ANR-10-INBS-07 PHENOMIN to YH]. The funders had no role in the study design, data collection and analysis, decision to publish, or preparation of the manuscript.

### Authors’ contributions

AR, YH, VB and EE designed the study and wrote the manuscript. AR and EE conducted the behavioural experiments and analysed the data. FdC provided the pipeline to analyse ultrasonic vocalisations. AR, CC and VN genotyped the animals.

## Acknowledgements

We would like to thank the members of the research group, of the IGBMC laboratory and of the ICS for their help, in particular Charley Pinault and Sophie Brignon for mouse breeding, and the people from the ICS Genotyping platform for genotype validation. We would also like to thank Nicolas Torquet for helpful comments on the manuscript.

