## Supplementary figures for "Day-to-day spontaneous social behaviours is quantitatively and qualitatively affected in a 16p11.2 deletion mouse model"

#### Supplementary Figure 1

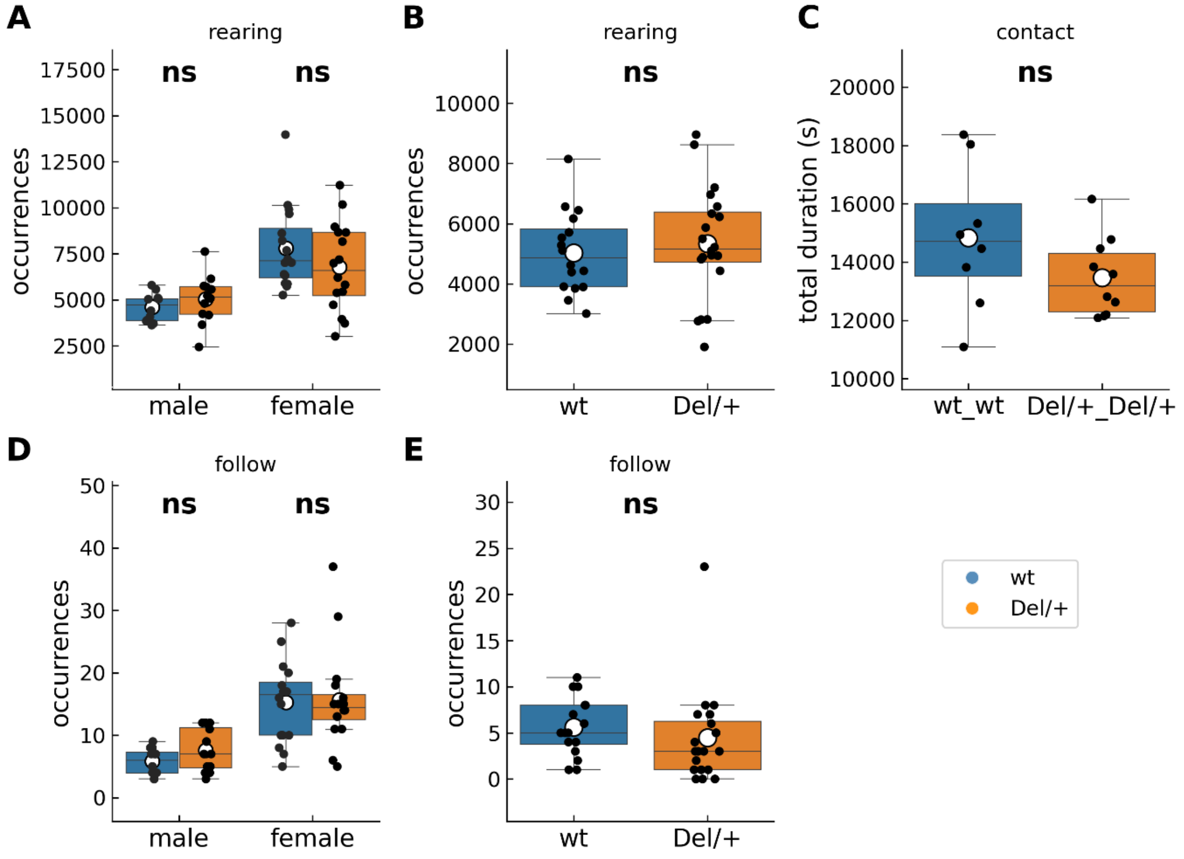


**Supplementary Figure 1: Vertical exploration and social behaviours displayed by mice of both sexes and genotypes in long-term monitoring**. (A) Total number of occurrences of rearings over the 3 nights of recording of spontaneous behaviours of mixed-genotype quartets of familiar males (12 wt and 12 Del/+ distributed in 6 cages) and females (16 wt and 16 Del/+ distributed in 8 cages). (B) Total number of occurrences of rearings over the 2 nights of recording of spontaneous behaviours of unfamiliar pairs of females of the same genotype (16 wt females distributed in 8 pairs and 20 Del/+ females distributed in 10 pairs). (C) Total time spent in contact over the 3 nights of recording of spontaneous behaviours of unfamiliar pairs of females of the same genotype (16 wt females distributed in 8 pairs and 20 Del/+ females distributed in 10 pairs). (D) Total number of occurrences of follow behaviours over the 3 nights of recording of spontaneous behaviours of mixed-genotype quartets of familiar males (12 wt and 12 Del/+ distributed in 6 cages) and females (16 wt and 16 Del/+ distributed in 8 cages). (E) Total number of occurrences of follow behaviours over the 2 nights of recording of spontaneous behaviours of unfamiliar pairs of females of the same genotype (16 wt females distributed in 8 pairs and 20 Del/+ females distributed in 10 pairs). (A, B, D, E) Linear Mixed Model, with genotype as fixed factor and cage/pair as a random factor; (C) Mann-Whitney U-test; ns: no significant effect of genotype, *: p<0.05, **: p<0.01, ***: p<0.001.

#### Supplementary Figure 2

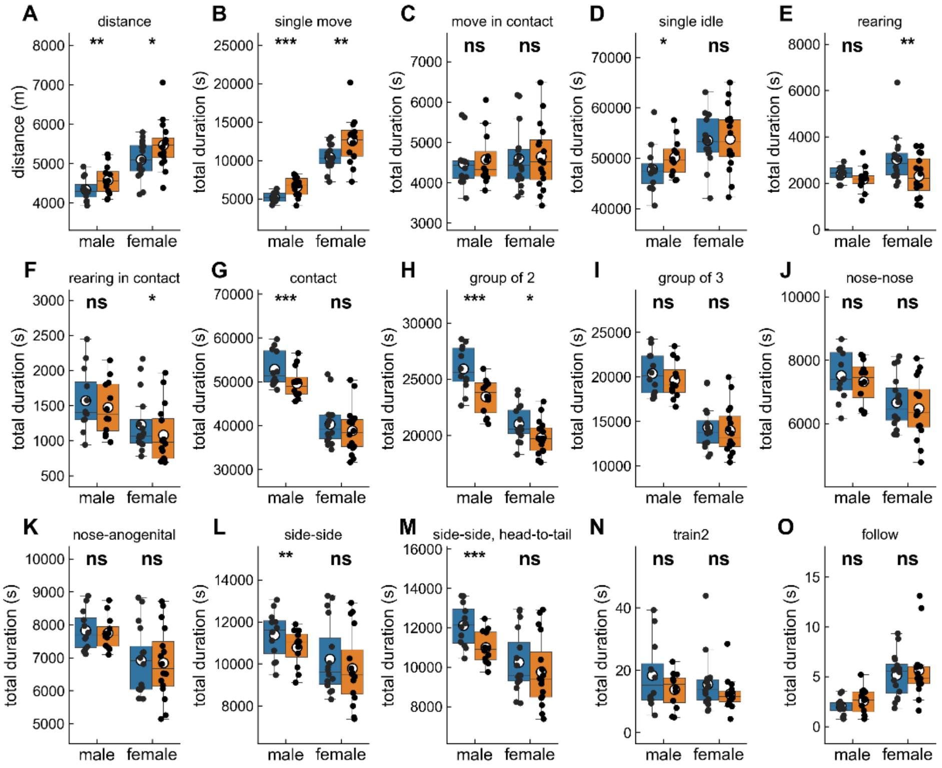


**Supplementary Figure 2: Raw data for the total duration of behavioural events displayed by mice of both sexes and genotypes in long-term monitoring**. (A) Distance travelled over the three nights of recordings of spontaneous behaviours of mixed-genotype quartets of familiar males (12 wt (blue) and 12 Del/+ (orange) distributed in 6 cages) or females (16 wt (blue) and 16 Del/+ (orange) distributed in 8 cages). (B-O) Total duration of behavioural events over the three nights of recordings of spontaneous behaviours of mixed-genotype quartets of familiar males (12 wt and 12 Del/+ distributed in 6 cages) or females (16 wt and 16 Del/+ distributed in 8 cages). Linear Mixed Model, with genotype as fixed factor and cage as a random factor: ns: no significant effect of genotype, *: $p<0.05$, **: $p<0.01$, ***: $p<0.001$.

### Supplementary Figure 3

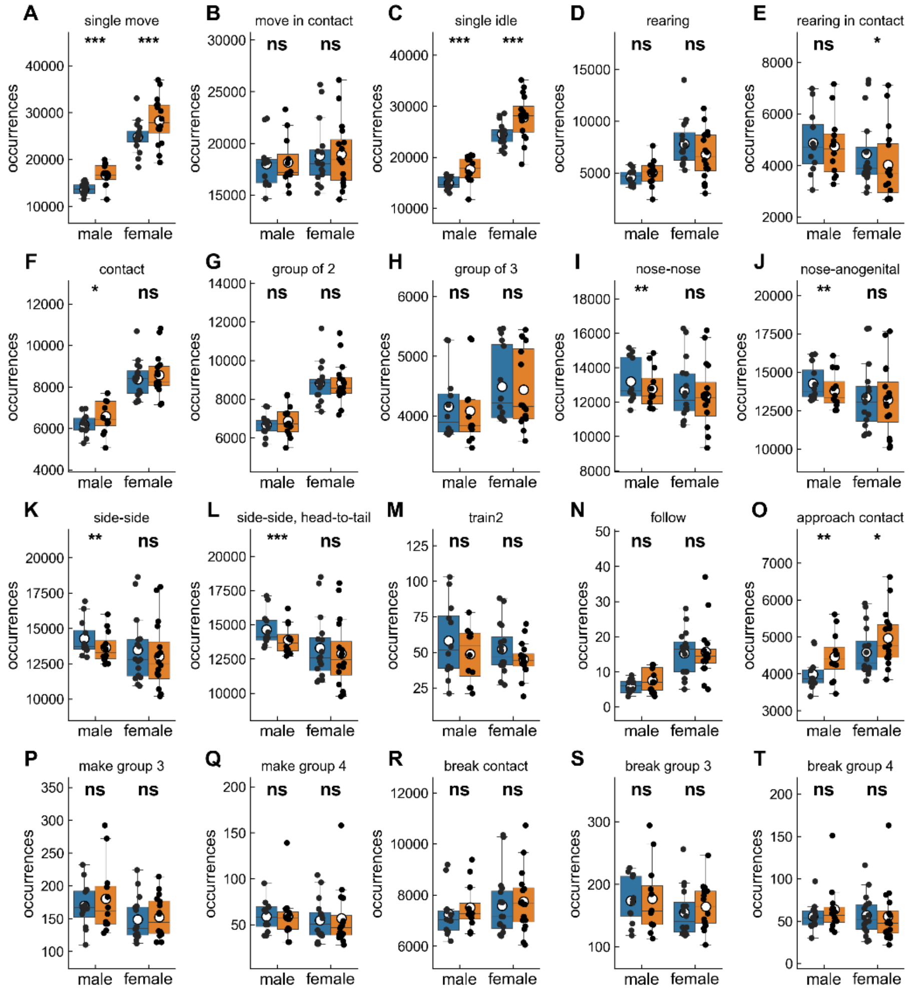


**Supplementary Figure 3: Raw data for the number of occurrences of behavioural events displayed by mice of both sexes and genotypes in long-term monitoring**. (A-T) Number of occurrences of behavioural events over the three nights of recordings of spontaneous behaviours of mixed-genotype quartets of familiar males (12 wt (blue) and 12 Del/+ (orange) distributed in 6 cages) or females (16 wt (blue) and 16 Del/+ (orange) distributed in 8 cages). Linear Mixed Model, with genotype as fixed factor and cage as a random factor: ns: no significant effect of genotype, *: $p<0.05$, **: $p<0.01$, ***: $p<0.001$.

### Supplementary Figure 4

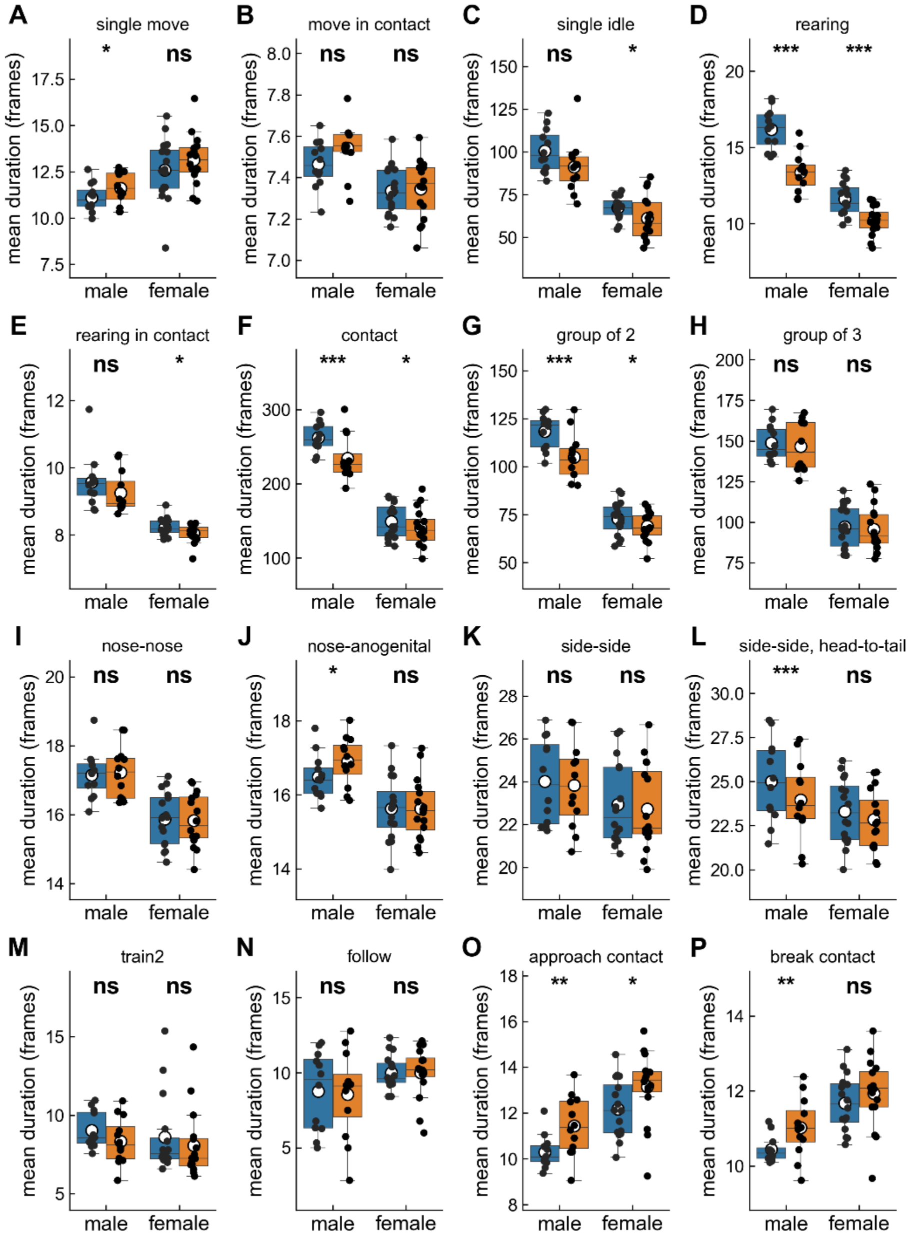


**Supplementary Figure 4: Raw data for the mean duration of behavioural events displayed by mice of both sexes and genotypes in long-term monitoring**. (A-P) Mean duration of behavioural events over the three nights of recordings of spontaneous behaviours of mixed-genotype quartets of familiar males (12 wt (blue) and 12 Del/+ (orange) distributed in 6 cages) or females (16 wt (blue) and 16 Del/+ (orange) distributed in 8 cages). Linear Mixed Model, with genotype as fixed factor and cage as a random factor: ns: no significant effect of genotype, *: p<0.05, **: p<0.01, ***: p<0.001.

### Supplementary Figure 5

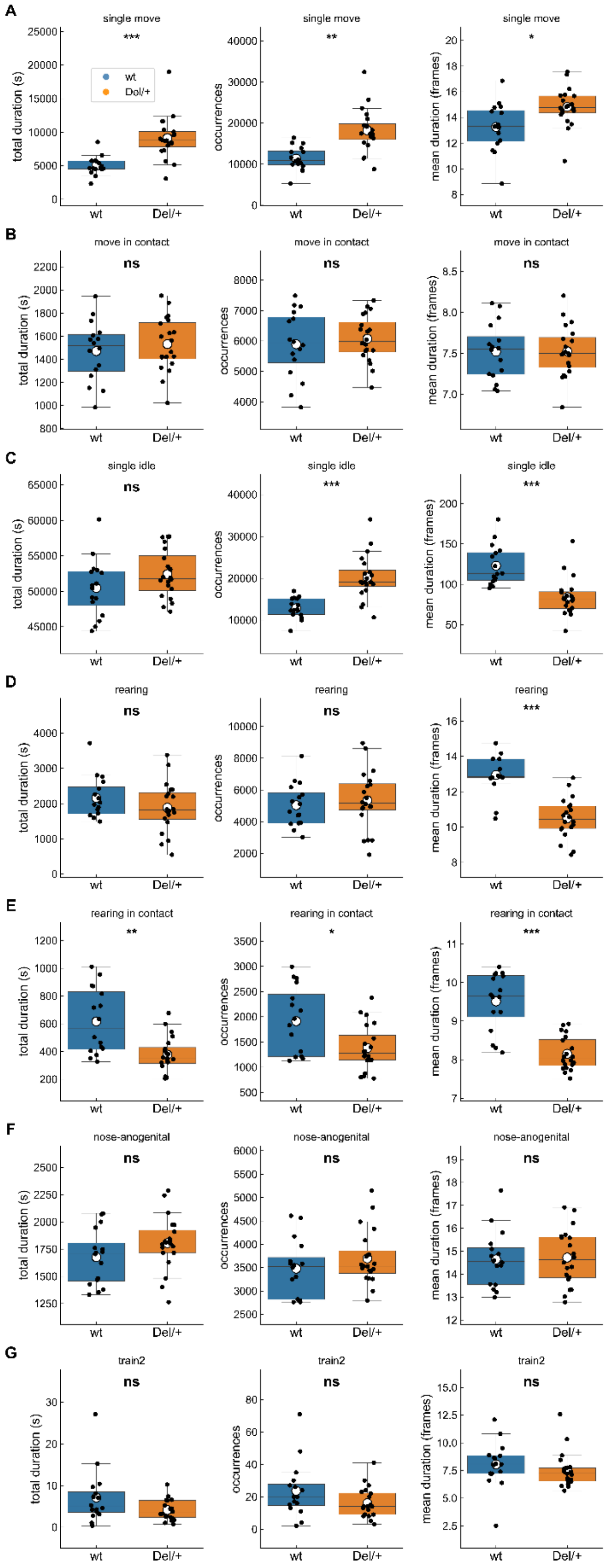

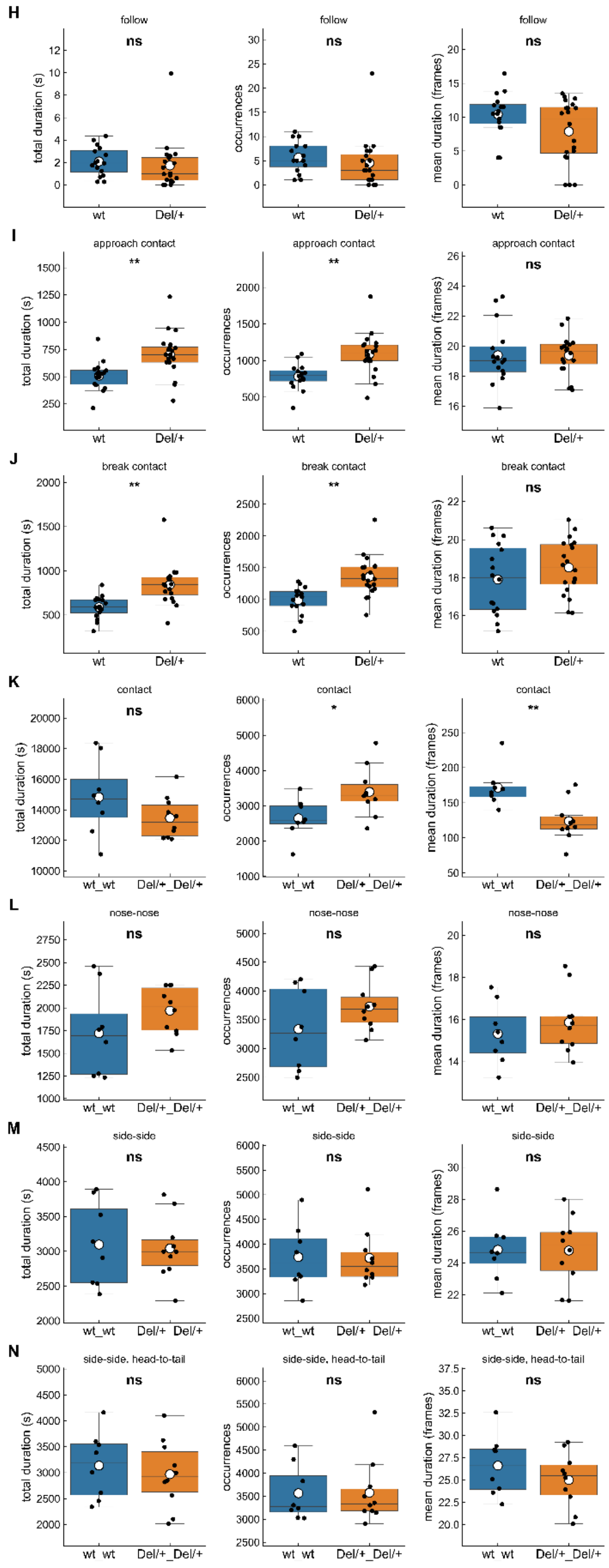


**Supplementary Figure 5: Raw data for the profile of unfamiliar pairs of females in long-term monitoring**. (A-P) Mean duration of behavioural events over the three nights of recordings of spontaneous behaviours of mixed-genotype quartets of familiar males (12 wt and 12 Del/+ distributed in 6 cages) or females (16 wt and 16 Del/+ distributed in 8 cages). Linear Mixed Model, with genotype as fixed factor and cage as a random factor: ns: no significant effect of genotype, *: p<0.05, **: p<0.01, ***: p<0.001.

### Supplementary Figure 6

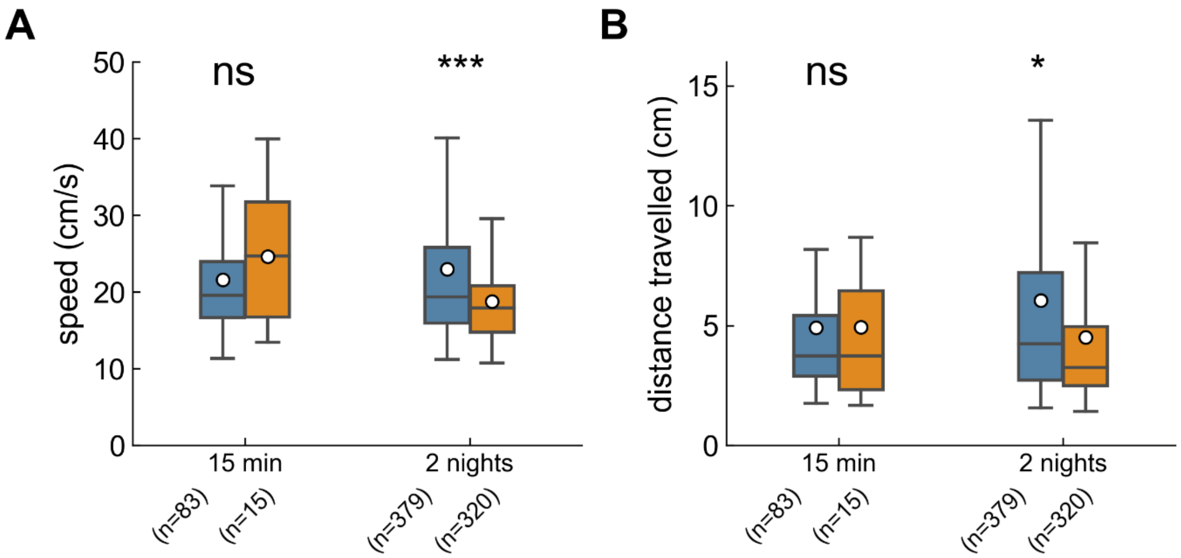


**Supplementary Figure 6: Characterisation of the follow behaviour with ano-genital sniffing (Train2) in the social encounters between same-genotype pairs of unfamiliar females**. (A) Mean speed of the animal during the Train2 events over short- and long-term recordings in wt-wt pairs (blue) and Del/+-Del/+ pairs (orange). (B) Mean distance travelled by the animal during the Train2 events over short- and long-term recordings in wt-wt pairs (blue) and Del/+-Del/+ pairs (orange). Linear Mixed Models, with genotype as a fixed factor and pair as a random factor; ns: no significant difference, *: $p<0.05$.

### Supplementary Figure 7

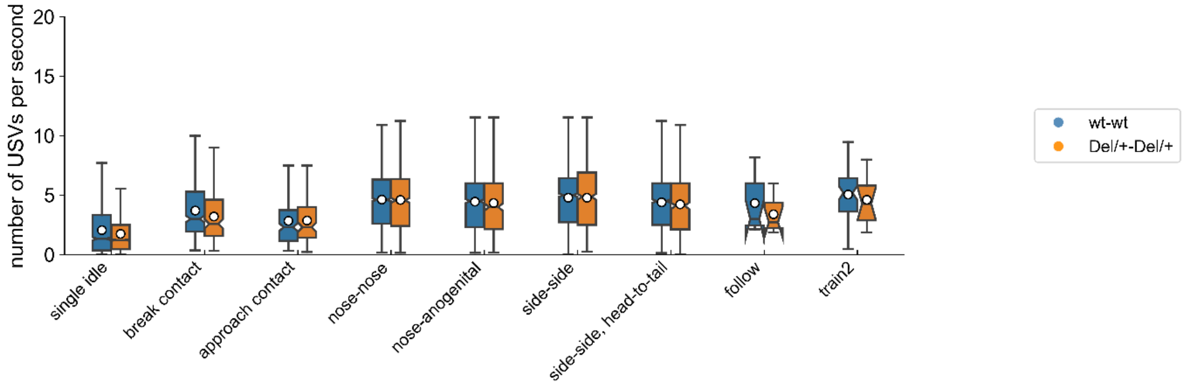


**Supplementary Figure 7: Rate of ultrasonic vocalisations emission during each behavioural event** during the social encounter between same-genotype pairs of unfamiliar females over long-term recordings. Linear Mixed Model with genotype as fixed factor and pair as random factor; ns: no significant difference.

### Supplementary Figure 8

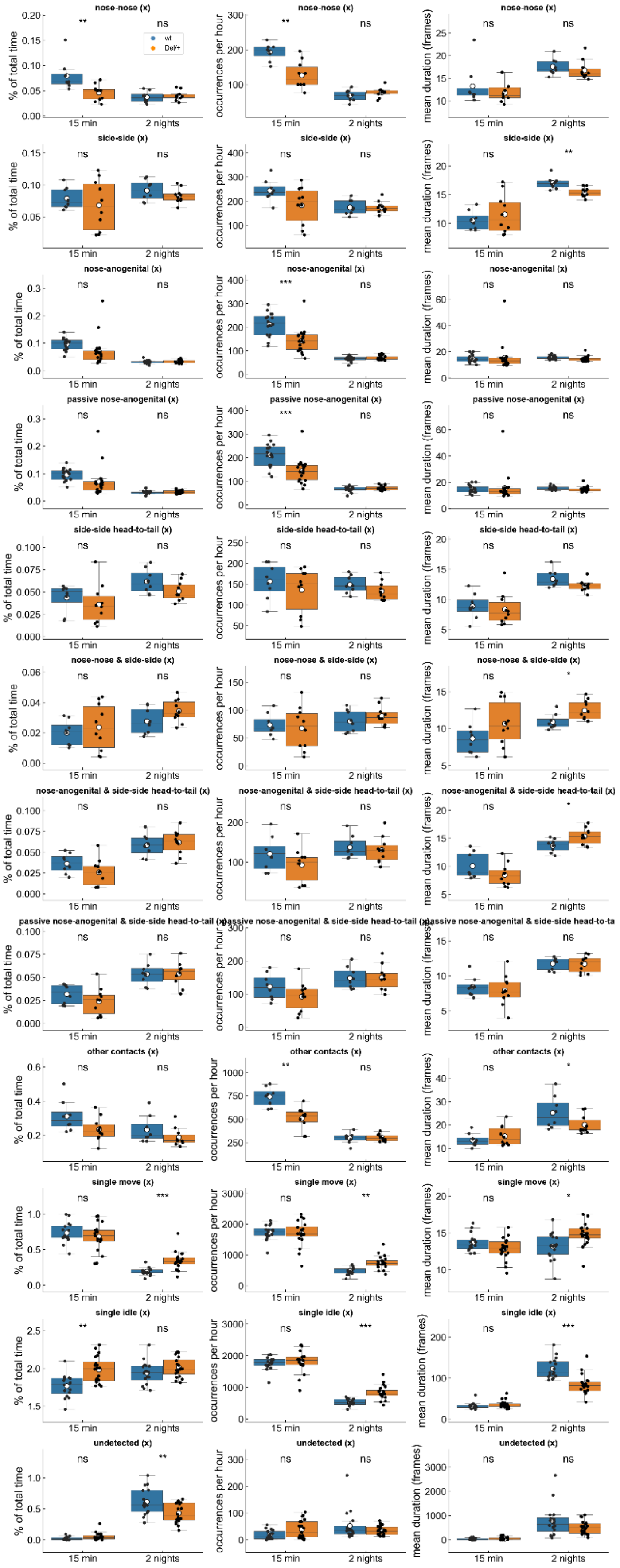

**Supplementary Figure 8: Behavioural profiles built with exclusive events in pairs of same-genotype unfamiliar females over short- (first 15 min) and long-term (2 nights) recordings**. Each row represents one exclusive behaviour, and for each exclusive behaviour the first column represents the proportion of total time spent in this behaviour, the second column represents the number of occurrences per hour and the third column represents the mean duration of this behaviour. 16 wt distributed in 8 pairs and 20 Del/+ distributed in 10 pairs; Linear Mixed Model or Mann-Whitney U-tests conducted; ns: no significant difference, *: $p<0.05$, **: $p<0.01$, ***: $p<0.001$.

### Supplementary Figure 9

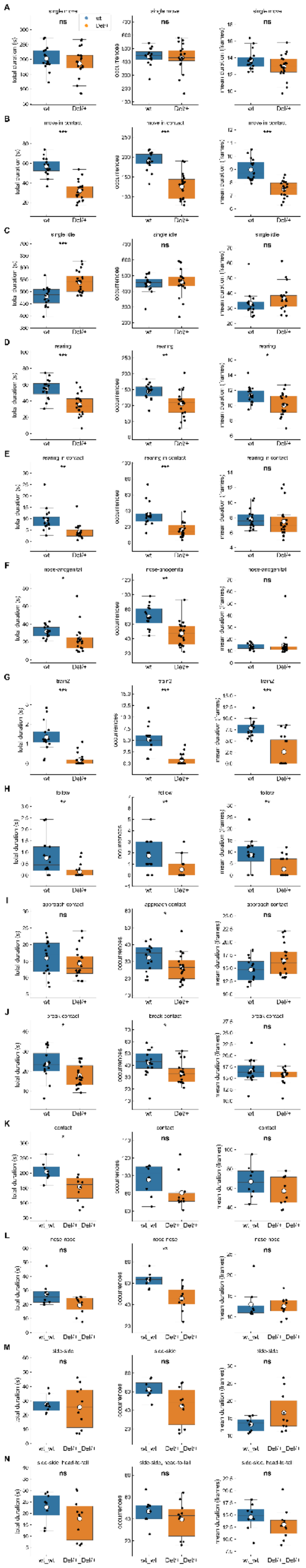

**Supplementary Figure 9: Raw data for the profile of unfamiliar pairs of females in short-term monitoring**. (A-P) total duration, number of occurrences and mean duration of behavioural events over the first 15 min of recordings of spontaneous behaviours of mixed-genotype quartets of familiar males (12 wt and 12 Del/+ distributed in 6 cages) or females (16 wt and 16 Del/+ distributed in 8 cages). Linear Mixed Model, with genotype as fixed factor and cage as a random factor: ns: no significant effect of genotype, *: $p<0.05$, **: $p<0.01$, ***: $p<0.001$.

### Supplementary Figure 10

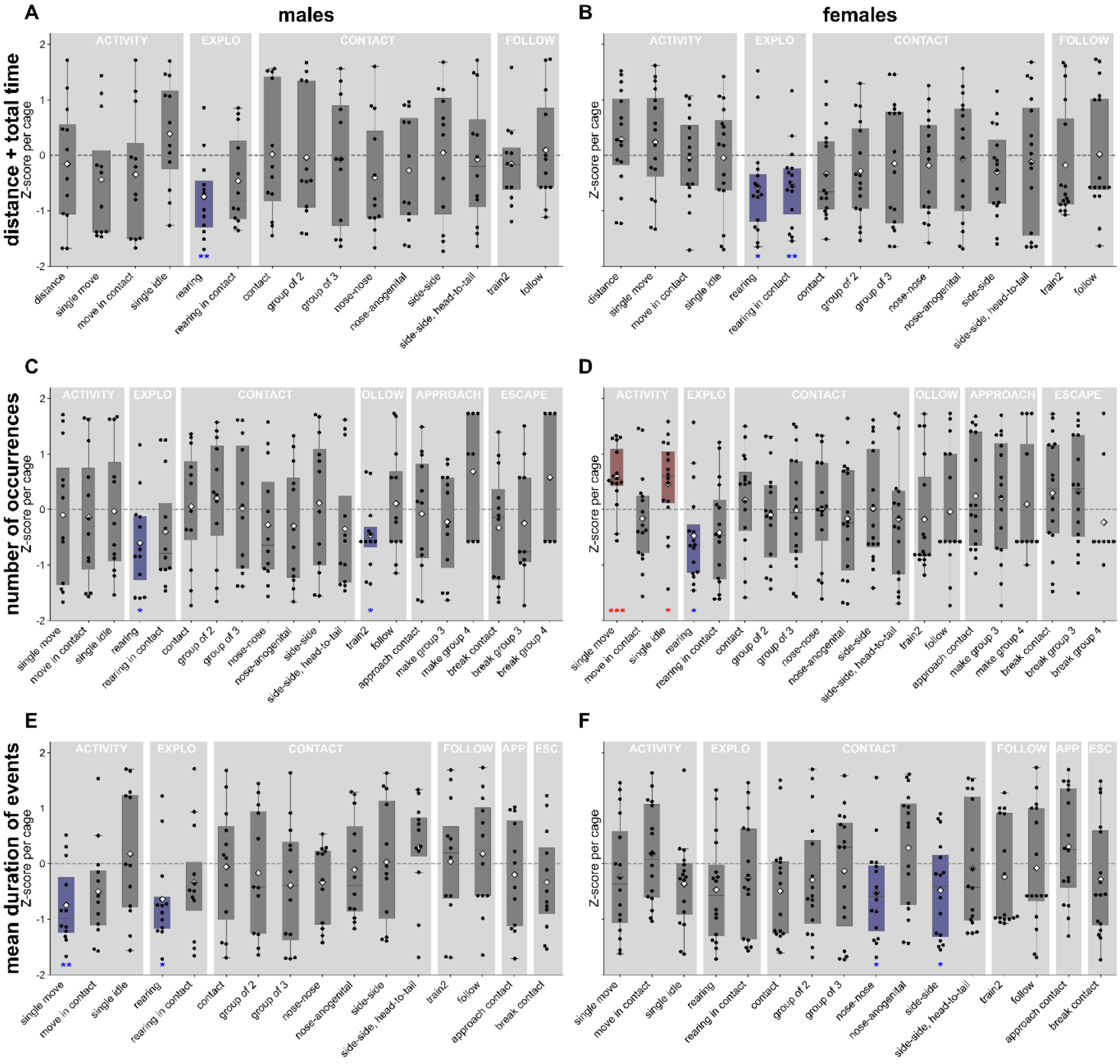


**Supplementary Figure 10: Behavioural profiles for the total duration, the number of occurrences and the mean duration of events of Del/+ males and females over the first 15 min in quartets.** (A, B) Z-score profile of the distance and the total duration of each behaviour for each Del/+ mouse compared to the mean behaviour of the four individuals within each quartet for males (A, n=12) as well as for females (B, n=16) over the first 15 min of recording. (C, D) Z-score profile of the number of occurrences of each behaviour for each Del/+ mouse compared to the mean behaviour of the four individuals within each quartet for males (C, n=12) as well as for females (D, n=16) over the first 15 min of recording. (E, F) Z-score profile of the mean duration of behavioural events for each Del/+ mouse compared to the mean value of the four individuals within each quartet for males (C, n=12) as well as for females (D, n=16) over the first 15 min of recordings. One-sample t-test if the data follow a normal distribution or Wilcoxon test if data are not normally distributed: *: p<0.05, **: p<0.01, ***: p<0.001. Red boxes and stars figure behavioural events that are more expressed in Del/+ mice compared to the mean of the whole cage; blue boxes and stars depict behavioural events that are less expressed in Del/+ mice compared to the mean of the whole cage; grey boxes reflect non-significant differences between Del/+ and the other animals of the cage.
